# Phylogenetic and protein structure analyses provide insight into the evolution and diversification of the CD36 domain ‘apex’ among scavenger receptor class B proteins across Eukarya

**DOI:** 10.1101/2023.11.13.566873

**Authors:** Reed T. Boohar, Lauren E. Vandepas, Nikki Traylor-Knowles, William E. Browne

## Abstract

The cluster of differentiation 36 (CD36) domain defines the characteristic ectodomain associated with scavenger receptor class B (SR-B) proteins. In bilaterians, SR-Bs play critical roles in diverse biological processes including innate immunity functions such as pathogen recognition and apoptotic cell clearance, as well as metabolic sensing associated with fatty acid uptake and cholesterol transport. While previous studies suggest this protein family is ancient, SR-B diversity across Eukarya has not been robustly characterized. We analyzed SR-B homologs identified from the genomes and transcriptomes of 165 diverse eukaryotic species. The presence of highly conserved amino acid motifs across major eukaryotic supergroups supports the presence of a SR-B homolog in the last eukaryotic common ancestor (LECA). Our comparative analyses of SR-B protein structure identifies the retention of a canonical asymmetric beta barrel tertiary structure within the CD36 ectodomain across Eukarya. We also identify multiple instances of independent lineage-specific sequence expansions in the apex region of the CD36 ectodomain — a region functionally associated with ligand-sensing. We hypothesize that a combination of both sequence expansion and structural variation in the CD36 apex region may reflect the evolution of SR-B ligand sensing specificity between diverse eukaryotic clades.

**Significance Statement:** SR-Bs are well-described in bilaterians, however the diversity and evolution of this ancient receptor protein family is poorly understood across Eukarya. Our analyses reveal a conserved beta barrel tertiary structure across eukaryotic SR-Bs that correlates with the presence of an intramolecular tunnel. In contrast, the putative ligand-sensing region associated with the apex of the protein can be highly divergent in both sequence and structure within and between taxa. Our data confirm the antiquity of the SR-B receptor protein family in Eukaryota and improve characterization of SR-B diversity outside of Metazoa. We hypothesize that eukaryotic SR-B diversity may be used as a model to explore receptor protein evolution driven by lineage-specific ligand specificity.

## Introduction

Cellular responses to external stimuli are often initiated by membrane-bound pattern recognition receptors (PRRs) (Koropatnick et al., 2004; Ausubel, 2005; Schaefer, 2014). Many PRR proteins contain functional domains associated with ligand binding and/or cellular adhesion, coupled with signal transduction (Wheeler et al., 2008; Özbek et al., 2010; Richter et al., 2018; Richter & Levin, 2019). Scavenger receptors (SRs) are a functionally defined group of structurally diverse transmembrane receptor proteins. Initially identified based on their function of ‘scavenging’ and removing modified lipoproteins, it has been established that SRs recognize a wide array of ligands and can serve as PRRs (reviewed in PrabhuDas et al., 2017; Taban et al., 2022).

Members of the class B scavenger receptors (SR-B), also commonly referred to as the ‘CD36 family’, are uniquely defined among SRs by two transmembrane domains with relatively short N- and C-terminal cytoplasmic tails involved in intracellular signal transduction and an ectodomain domain constituting the CD36 antigen (Taban et al., 2022; Figure 1A). Within mammals, several multifunctional class B scavenger receptors have been described: SCARB1, lysosomal integral membrane protein type 2 (LIMP-2 or SCARB2), and CD36 (also known as SCARB3). In vertebrates, SR-B proteins are expressed in a variety of cell types and bind diverse ligands including lipid-based pheromones, lipid-soluble vitamins, cell adhesion proteins, cholesterols, phospholipids, microbe-associated molecular patterns (MAMPs), as well as damage-associated molecular patterns (DAMPs) from cell debris, and long-chain fatty acids (LCFAs) (Koropatnick et al., 2004; Ausubel, 2005; Silverstein et al., 2009; Sheedy et al., 2013; Schaefer, 2014; Chen et al., 2022). In mammalian microglial cells, dysregulation of CD36 may be involved in the pathologies of Alzheimer’s Disease (Stuart et al. 2007; Park et al., 2013; Dobri et al., 2021). SR-B proteins that recognize MAMPs and DAMPs also cooperate with other cell surface PRRs to mediate innate immune responses including phagosome/endosome formation (Erdman et al., 2009; Silverstein et al., 2009; Stewart et al., 2010; Chen et al., 2022).

**Figure 1:**
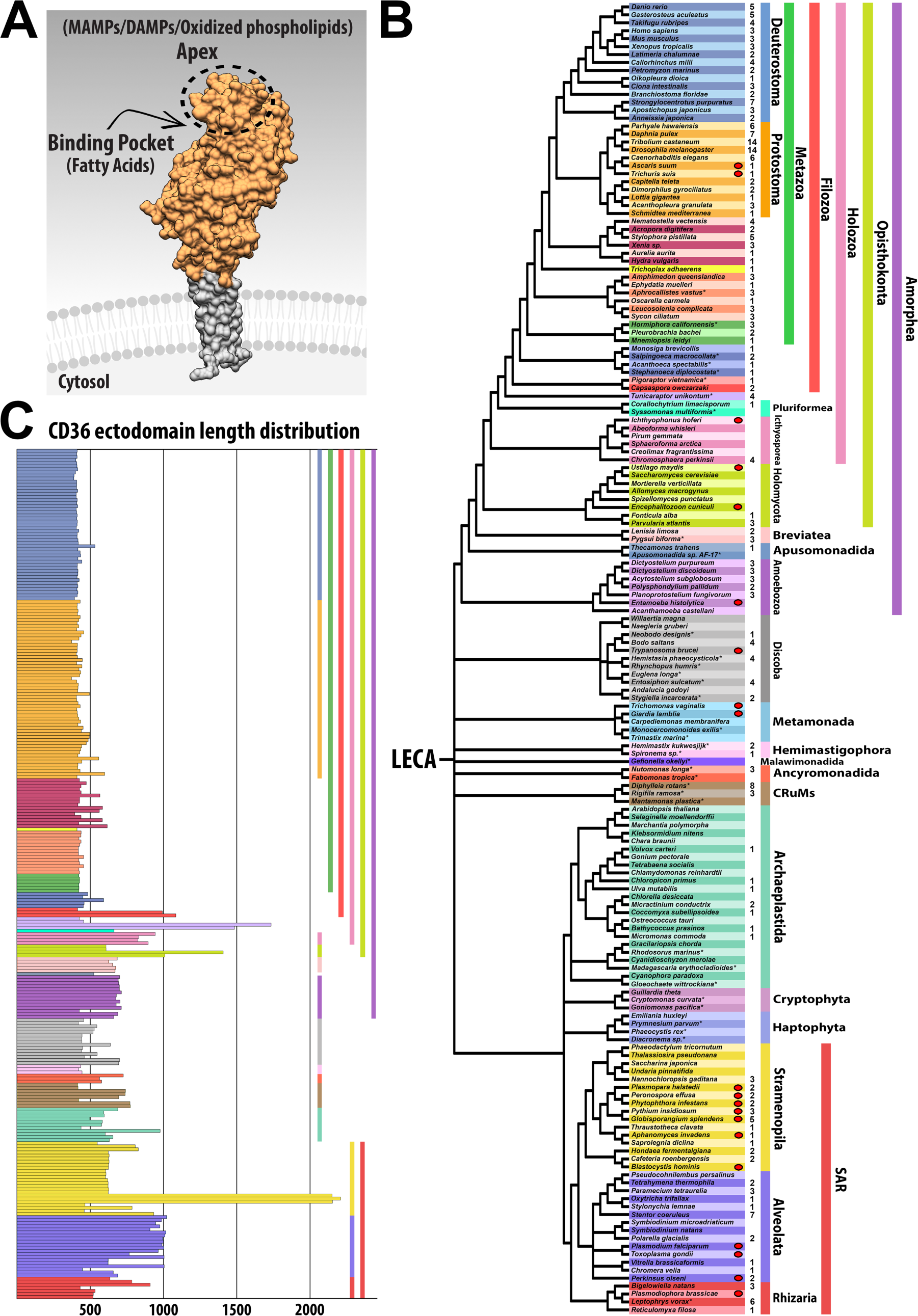
Distribution of SR-Bs across Eukarya. **A)** Space-filling model of SCARB3. SR-Bs are integral membrane proteins with two transmembrane domains, cytoplasmic N- and C-terminal tails (grey shading) and a CD36 ectodomain (orange shading) characterized by a ligand sensing apex, binding pocket and containing an intramolecular tunnel. **B)** Phylogenetic distribution of SR-Bs. Asterisk indicates taxa for which transcriptomic data was used. A red oval next to a row indicates obligate parasitic life history. Number indicates how many SR-B homologs were detected in each representative taxon. Color-coded taxonomic groupings are on the right. Phylogeny based on Burki et al. (2020), Carr et al. (2017), Derelle et al. (2016), Ettahi et al. (2021), Li X et al. (2021), Li Y et al. (2021), Strassert et al. (2021). Raw CD36 Pfam domain hits following isoform removal for each taxon can be found in Supplementary Table 2. **C)** Length distribution for each of the 279 CD36 ectodomains. Includes amino acids comprising the predicted non-cytoplasmic region containing CD36 ectodomain sequence between transmembrane helices. Colors follow the taxonomic scheme in panel (B).

X-ray crystallography of human SR-B proteins shows that the CD36 ectodomain folds into an asymmetric beta barrel structure with a group of alpha helices at the protein’s ‘apex’ – the most distal region of the ectodomain extending farthest from the cell membrane (Neculai et al., 2013; Hsieh et al 2016; Supplementary Figure 1). This “three-helix bundle” at the apex plays a critical role in ligand recognition (Kar et al., 2008; Kuda et al., 2013; Neculai et al., 2013; Supplementary Figure 1A). The apex region is spatially adjacent to a binding pocket and an intramolecular tunnel that assists with transport of cholesterol esters and LCFAs (Figure 1A; Neculai et al., 2013; Conrad et al., 2017; Glatz et al., 2018; Heybrock et al., 2019). Additionally, the ectodomain contains 2-3 disulfide bridges between loop regions of the asymmetric beta barrel core (Papale et al., 2011; Yu et al., 2012). SR-B proteins have been identified and functionally characterized in diverse metazoans, from mammals to sponges (Franc et al., 1996; Müller et al., 2004; Dinguirard & Yoshino, 2006; Means et al., 2009; Bi et al., 2015; Neubauer et al., 2016). Insects have three distinct lineage-specific clusters of CD36 domain-containing proteins (Nichols et al., 2008) with similar structural features to vertebrate homologs, including the three-helix bundle at the apex of the ectodomain, a putative binding pocket, presence of an intramolecular tunnel, and disulfide bridges (Nichols and Vogt, 2008; Gomez-Diaz et al., 2016). Intriguingly, the amoebozoan slime mold *Dictyostelium discoideum* possesses three LIMP-2/CD36 genes (LmpA-C) that have been shown to function in phagocytosis (Karakesisoglou et al., 1999, Sattler et al., 2018). However, little is known regarding the distribution of SR-Bs or their diversity and evolution in other non-metazoan eukaryotes.

To investigate SR-Bs across a diversity of eukaryotic lineages, we leveraged comparative analysis of proteins using sequence alignments, amino acid motif prediction (Bailey, 2021), and AI algorithm tools for both protein structure prediction (Baek et al., 2021; Jumper et al., 2021) and small molecule interaction prediction (Hekkelman et al., 2023). Our phylogenetic analyses support the presence of an ancestral SR-B-like protein in the Last Eukaryote Common Ancestor (LECA). Protein structure predictions show that eukaryotic SR-Bs share a broadly conserved asymmetric beta barrel tertiary structure within the CD36 ectodomain that contrasts with an evolutionarily labile region at the membrane-distal apex of the ectodomain. This membrane-distal apex region correlates with lineage-specific sequence expansions of highly variable alpha helical structure. Importantly, in functionally characterized SR-Bs, amino acid residues critical for ligand recognition reside in the apex region (Neculai et al., 2013; Hsieh et al., 2016; Supplementary Figure 1A). Based on our results, we hypothesize that sequence expansion and structural divergence in the apex region of the CD36 ectodomain may reflect the evolution of SR-B ligand specificity between diverse eukaryotic lineages.

## Results

### SR-B-like proteins are present across diverse eukaryotes and characterized by lineage-specific sequence expansions

SR-B protein sequences are operationally defined as possessing a CD36 ectodomain flanked by two transmembrane helices (PrabhuDas et al., 2017; this study). We used HMMER to search for the presence of CD36 domains in 165 publicly available eukaryotic proteomes and recovered a total of 418 unique CD36 domain-containing protein sequences (Supplementary Table 1; Supplementary Table 2). CD36 domain-containing protein sequences were then screened for flanking transmembrane domains, from which we identified 279 *bona fide* SR-B homologs. SR-B homologs were recovered across Eukarya, including in taxa from several early-diverging eukaryotic lineages including the Hemimastigophora, Ancyromonadida and CRuMs (Figure 1B).

Within Amorphea, SR-B homologs were detected in every major clade, including Metazoa (143), Choanoflagellata (5), Ichthyosporea (4), Holomycota (4), and Amoebozoa (14). We also identified SR-Bs in the transcriptome of *Tunicaraptor unikontum* (4), the pluriformean *Corallochytrium limacisporum* (1), the Breviatea (5), and the apusomonadidan *Thecamonas trahens* (1). Notably, within Ichthyosporea and Holomycota, *bona fide* SR-B homologs were only recovered from early-diverging lineages, suggesting loss of SR-Bs in more derived taxa (Figure 1B; Supplementary Table 2).

SR-B homologs were detected in seven chlorophyte taxa within Archaeplastida; however, no SR-B homologs were recovered from streptophytes, glaucophytes, or rhodophytes (Figure 1B; Supplementary Table 2), suggesting independent losses in each of these lineages. We identified SR-B homologs in each of the three major clades that constitute the SAR supergroup: <u>S</u>tramenopila (24), <u>A</u>lveolata (20), and <u>R</u>hizaria (10). Within Stramenopila, all sampled oomycetes possess SR-B homologs. Among Alveolata, all representative ciliates possessed at least one SR-B homolog, except for the fish parasite *Pseudocohnilembus persalinus*. Eight SR-Bs, the largest SR-B complement in a non-metazoan taxon, were detected in *Diphylleia rotans*, a member of the CRuMs lineage. Among early-diverging eukaryotic lineages we detected the following complements of SR-B homologs: Discoba (15), Hemimastigophora (3), Ancyromonadida (3), and CRuMs (11).

Some recovered SR-B protein sequences with unusually long cytoplasmic regions appeared to represent gene model fusions and contained protein domains not typically associated with SR-Bs. For example, the ctenophore *Mnemiopsis leidyi* SR-B gene (ML01096a) contains a long N-terminal cytoplasmic region with a PHD finger motif that is not found in other eukaryotic SR-Bs. We compared transcript data to the genomic locus and identified exons 1–7 as belonging to a separate PHD finger-containing gene, while exons 8 and 9 of the gene model correspond to SR-B transcripts, with a putative transcription start site 75 bp upstream of exon 8 (Moreland et al., 2014). A revised protein sequence translated from this start codon through exons 8 and 9 was used in this study (Supplementary File 1). Similarly, the choanoflagellate *Monosiga brevicollis* sequence (XP_001747074.1) also likely represents a gene model fusion based upon comparison with other choanoflagellate SR-B transcripts (Richter et al., 2018). A revised protein sequence, translated from exons 2–11 was used in this study (Supplementary File 1).

We then isolated and aligned CD36 ectodomain amino acid sequences from recovered *bona fide* SR-B homologs by removing predicted sequences for transmembrane helices and cytoplasmic tails (Supplementary File 2). We recovered significant lineage-specific variation in sequence length among CD36 ectodomains across eukaryotes. Metazoan CD36 ectodomains are between 367aa and 616aa, whereas non-metazoan CD36 ectodomains vary more widely, between 413aa and 2294aa (Figure 1C). In all cases, the most substantial length variation is associated with the membrane-distal apex of the ectodomain (Supplementary File 3, positions 433–2064). Notably, this variable sequence expansion region corresponds with two alpha helices of the three-helix bundle that is known to play a functional role in mediating ligand recognition in mammalian SR-Bs (Neculai et al., 2013).

### Phylogenetic analysis of eukaryotic SR-B proteins

We employed both maximum likelihood (RAxML) and Bayesian (Mr. Bayes) phylogenetic analyses to explore relationships between CD36 domains of SR-B homologs across Eukarya. Congruent bootstrap (BS ≥ 80%) and Bayesian posterior probability (BPP ≥ 0.9) support were recovered for shallow nodes that unite SR-B orthologs and identify putative lineage-specific paralogs, including some informative clades among metazoans (Figure 2; Supplementary Figure 2; Supplementary Figure 3; Supplementary Table 3). However, results from both analyses are broadly typified by relatively long branch lengths and a lack of support for deep divergences. For example, vertebrate SCARB2 orthologs group with high support (BS = 100%, BPP = 1) indicating an ancestral SCARB2 protein was likely present in the craniates. However, despite support for each of the three insect SR-B groups (Nichols and Vogt, 2008), the relationships between emp-like, crq-like, and SNMP-like insect SR-Bs to other metazoan SR-Bs remain unresolved (Figure 2; Supplementary Table 3).

**Figure 2:**
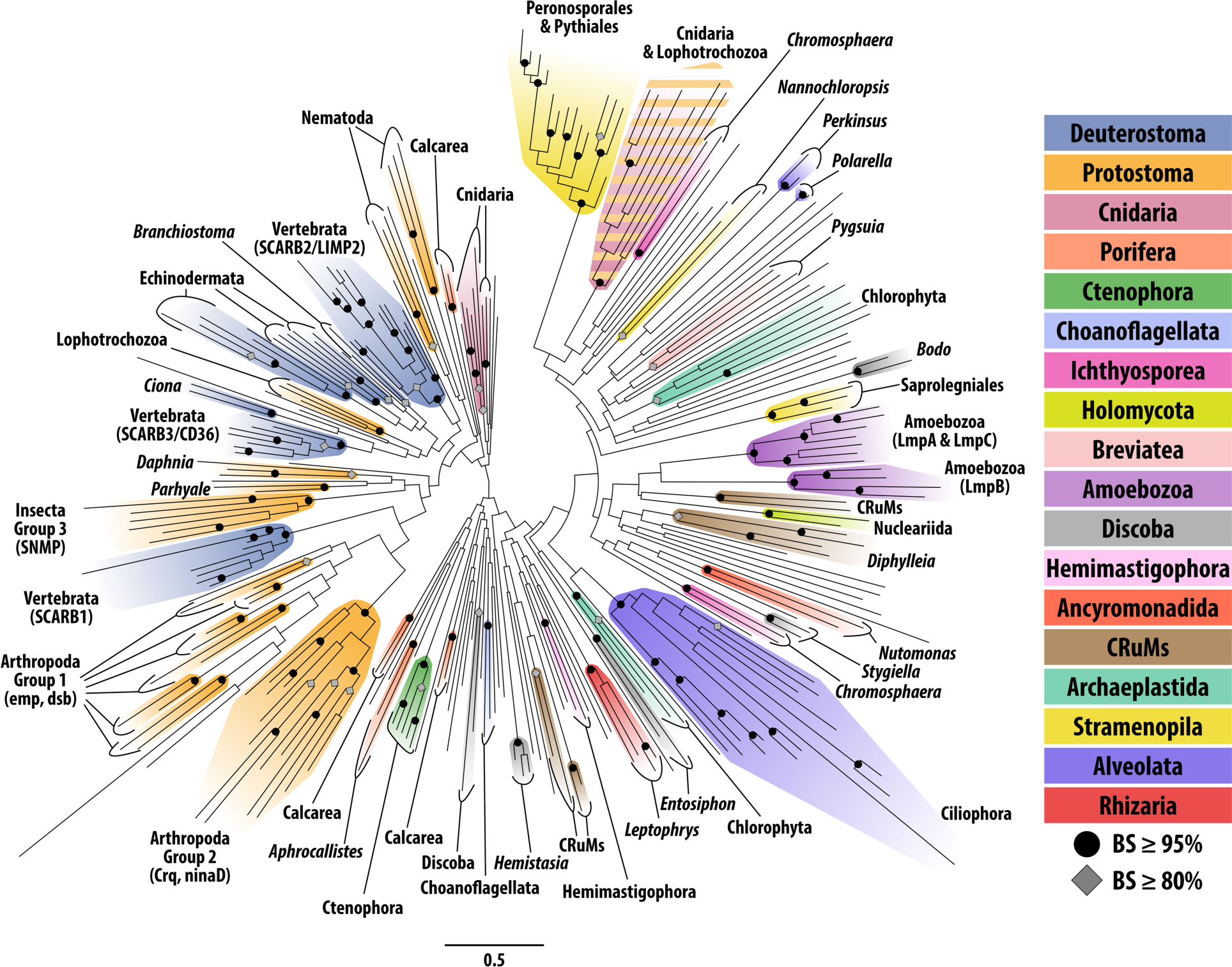
Maximum likelihood (RAxML) gene tree for eukaryotic CD36 domains. Black circles indicate nodes with bootstrap support ≥ 95%. Gray diamonds indicate nodes with bootstrap support ≥ 80%. Clades with ≥ 80% bootstrap support are colored according to Figure 1B. Unannotated tree with node support values is Supplementary Figure 2.

We did not recover support for deep relationships between non-metazoan SR-B homologs. For example, the largest supported clade of non-metazoan SR-Bs is restricted to Peronosporales and Pythiales oomycetes. Among amoebozoans, SR-Bs clustered into two strongly supported clades, one containing putative LmpB-like homologs and the other including both LmpA-like and LmpC-like homologs (Figure 2; Supplementary Table 3; Sattler et al., 2018). Notably, all clades united by nodes with BS ≥ 80% were composed of SR-Bs from monophyletic taxa, with the exception of a single clade comprising six cnidarian and two lophotrochozoan sequences (Figure 2; Supplementary Table 3). In contrast to the majority of metazoan SR-Bs with CD36 ectodomains <500 aa, the cnidarian and lophotrochozoan CD36 ectodomains in this group are all >500 aa in length (Figure 1C). While these eight sequences share three invariant residues (F788, W812, and L1188; Supplementary File 3) and an additional independant alignment identified a further three shared invariant residues (Supplementary Figure 4), we are unable to reject the possibility of long branch attraction.

Our combined data suggests that SR-Bs appear to have undergone significant sequence divergence during the evolution and diversification of eukaryotic lineages, obscuring homolog relationships between lineages. Additionally, eukaryotic genomic diversity remains underrepresented despite significant recent progress in the availability of genomic resources among non-metazoans (Richter et al., 2022). In this study ∼45% of CD36 domain-containing sequences are metazoan, thus taxonomic sampling bias represents a potential limitation for resolving phylogenetic relationships among SR-B homologs. Despite these limitations, we interpret our phylogeny reconstruction as supporting the presence of at least one ancestral SR-B in LECA (Figure 2: Supplementary Figure 2: Supplementary Figure 3).

### CD36 disulfide bridge conservation in Metazoa

Three pairs of cysteine residues forming intramolecular disulfide bridges have been identified in the CD36 ectodomain of several mammalian and insect SR-Bs (Supplementary Figure 1A; Rasmussen et al., 1998). Functional analyses have highlighted the importance of these disulfide bridge-forming cysteines for primate SCARB1 and SCARB3 multimerization, as well high-density lipoprotein (HDL) binding and HDL cholesterol ester uptake by SCARB1 in primates (Thorne et al., 1997; Papale et al., 2011; Yu et al., 2012). However, binding of oxidized low-density lipoproteins (oxLDLs) by human SCARB3 appears to be independent of disulfide bridge formation (Jay et al., 2015). We characterized the distribution of cysteine residue pairs with putative homology to human SCARB3 disulfide bridges-1, -2, and -3 across eukaryotic SR-Bs (Supplementary Table 4).

Disulfide bridge-1 and disulfide bridge-2 cysteine residue pairs are well-represented among bilaterian SR-B sequences (83%) and both disulfide bridge pairs are detected among cnidarian, placozoan, and sponge sequences. Cysteine residue pairs with putative homology to disulfide bridge-3 appear to be specific to bilaterians, and among vertebrates are restricted to SCARB3 homologs (Supplementary Table 4). Among protostomes, disulfide bridge-3 cysteines are found in most arthropod sequences (90%). However, disulfide bridge-3 cysteines appear to be broadly absent from representative lophotrochozoan sequences, only being recovered from the annelid *Dimorphilus gyrociliatus*. Cnidarian and lophotrochozoan CD36 ectodomains >500 aa in length appear to lack homologous cysteine residues. Notably, no putative disulfide bridge-forming cysteine homolog pairs were identified in representative ctenophore CD36 ectodomain sequences.

Among non-metazoan sequences, the only cysteine pairs recovered were from three saprolegnialean oomycete SR-Bs (Thrcl_SR-B, Aphin_SR-B, Sapdi_SR-B; Supplementary Table 4). However, upon visual inspection, these putative disulfide bridge-3 cysteines were found to belong to a region of lineage-specific sequence expansion (Supplementary File 2) and thus excluded as false positives. Therefore, among representative non-metazoans no putative disulfide bridge-forming cysteine pairs were detected in our analysis.

### Motifs within the CD36 ectodomain

To further characterize patterns of sequence conservation and lineage-specific divergence in the CD36 ectodomain across eukaryotic SR-Bs, we performed an amino acid motif discovery analysis using the STREME algorithm to identify enriched motifs (Bailey, 2021). We recovered eighteen motifs that broadly characterize the CD36 ectodomain across the 279 identified eukaryotic SR-B sequences (Figure 3A; Supplementary Figure 5). Conserved regions were associated with high significance motifs; in contrast, lineage specific regions with lower conservation were associated with relatively lower motif significance (Supplementary Table 5). Motif position weight matrices were used to probe individual SR-B sequences to predict motif presence (Figure 3B).

**Figure 3:**
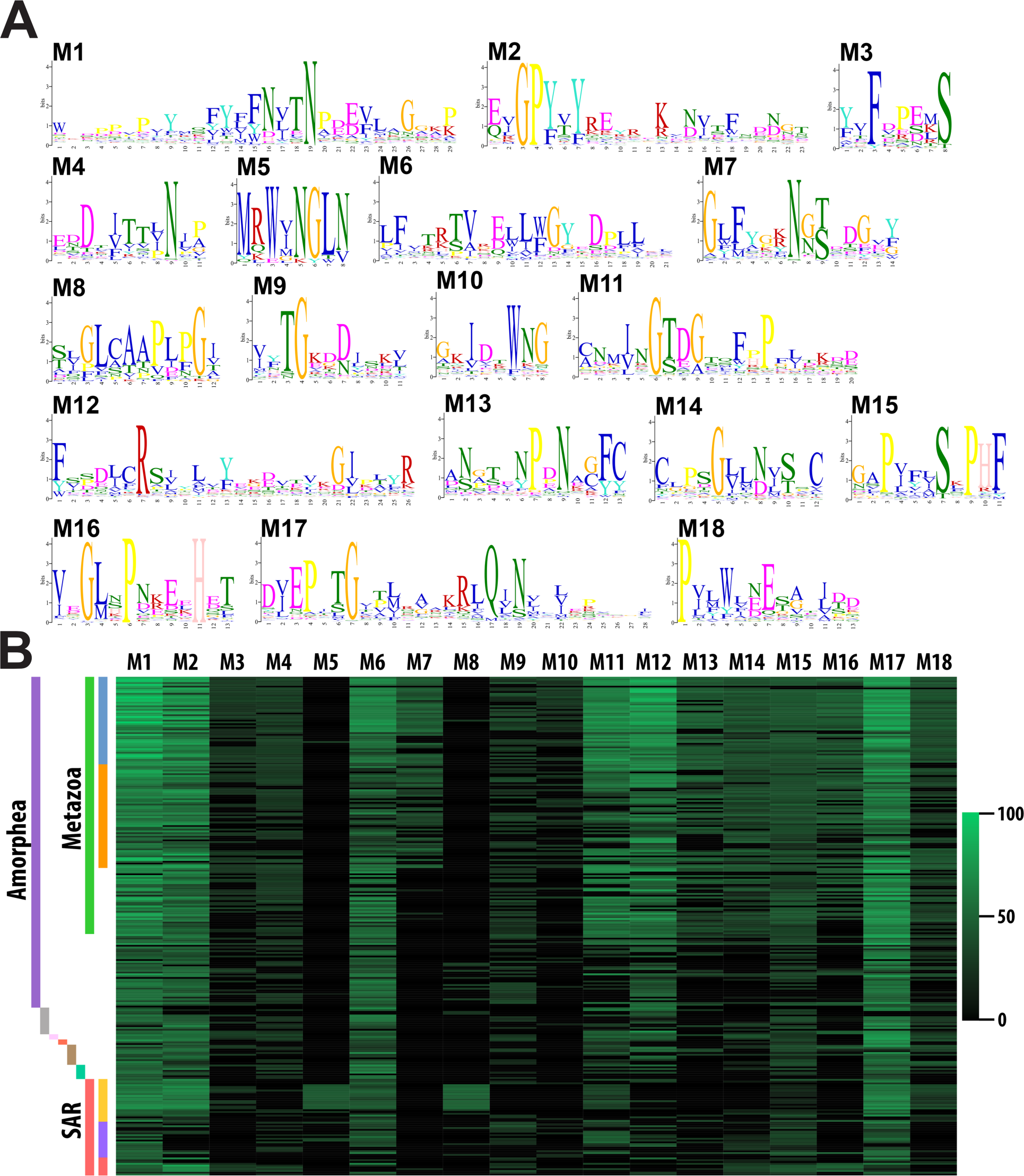
CD36 ectodomain amino acid motifs. **A)** Sequence logos for 18 amino acid motifs recovered from our set of 279 eukaryotic CD36 domains by STREME v5.5.3. Letter height represents the relative amino acid frequency. **B)** A heatmap depicting the sequence score for each motif to each eukaryotic CD36 ectodomain is shown. Raw sequence scores calculated by STREME were transformed to a 0–100 scale represented on the right. The full matrix of motif sequence scores is found in Supplementary Table 5. Colors correspond with the taxonomic scheme in Figure 1B.

Motifs M1, M2, and M17 are detected with high confidence in >90% of recovered SR-B sequences (Supplementary Table 5). These three motifs correspond with the most highly conserved regions of the unfiltered alignment (Supplementary File 3). Motifs M1 and M2 are located in the N-terminal region and motif M17 is located in the C-terminal region of the CD36 ectodomain. These dominant motifs contribute structurally to the longest β-strands within the conserved asymmetric beta barrel of the CD36 ectodomain and are spatially adjacent (Supplementary Figure 5; Figure 4C; Figure 5C). Motif M6 is also highly represented across eukaryotes (211 out of 279 sequences) and corresponds with a relatively short β-strand and α-helical region linking two α-helices in the apex (Supplementary Figure 5; Supplementary Figure 8).

**Figure 4:**
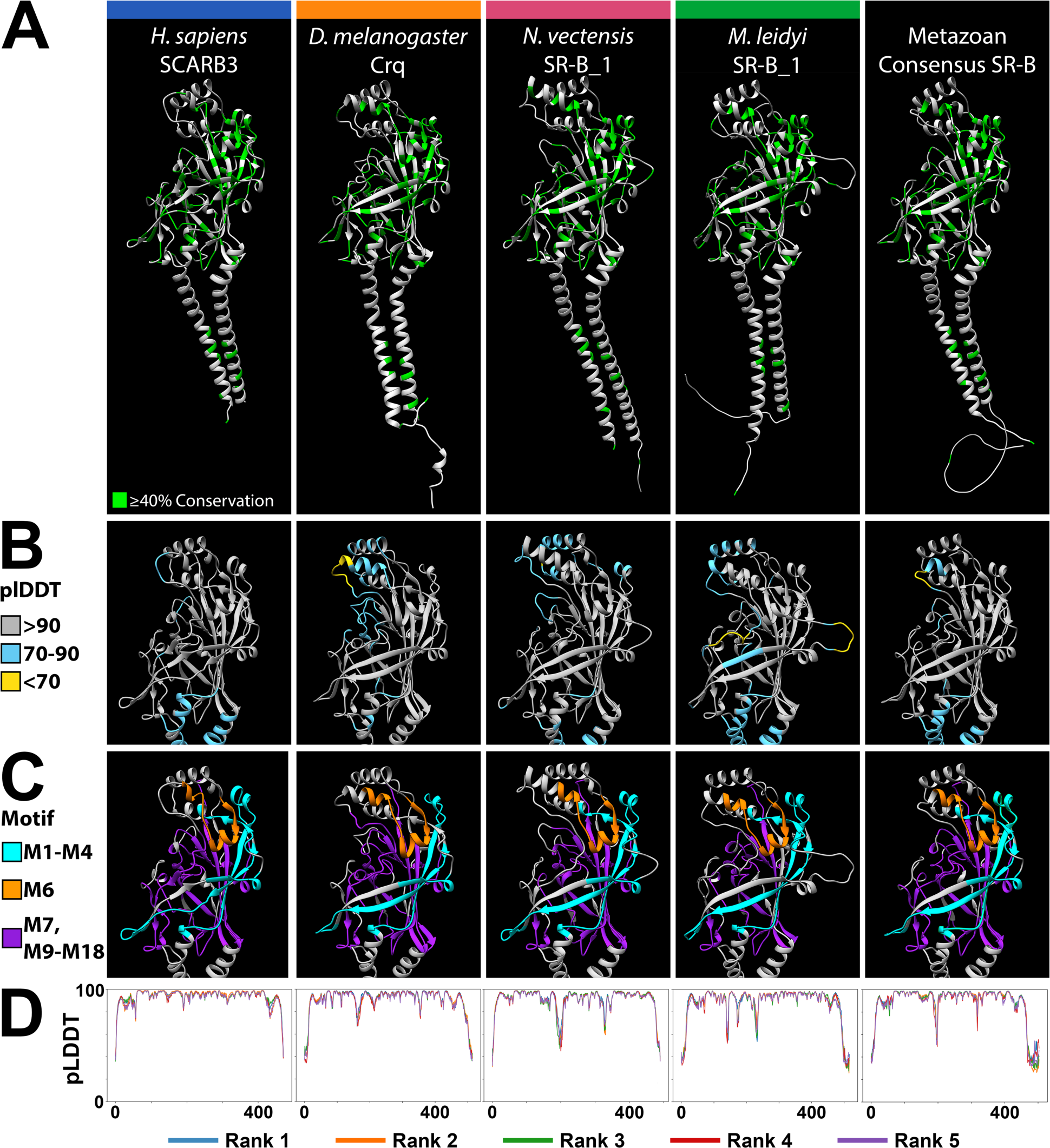
Metazoan SR-Bs share a highly conserved tertiary structure. The highest ranked AlphaFold2 structure predictions for four exemplar metazoan SR-B proteins. From left to right, *H. sapiens* SCARB3, *D. melanogaster* Crq, *N. vectensis* SR-B_1, and *M. leidyi* SR-B_1, alongside the consensus structure prediction derived from an all-metazoan SR-B sequence alignment. Protein structure visualized with Chimera. **A)** Green residues in the ribbon models reflect ≥40% sequence conservation across eukaryotic SR-Bs. **B)** pLDDT scores (predicted Local Distance Difference Test) computed by AlphaFold2 assess per-residue confidence across the CD36 ectodomain region of each model. Residue color reflects high confidence (gray), moderate confidence (light blue), and low confidence (yellow) pLDDT (Varadi et al., 2021). **C)** The CD36 ectodomain region of each model is annotated with the amino acid motifs discovered via STREME. Motifs M1–M4 (cyan) lie more N-terminal to the three-helix bundle at the apex, and motifs M7, M9–M18 (purple) lie more C-terminal to the three-helix bundle at the apex. Motif M6 (orange) connects α5 to α7 but is not part of the bundle itself. **D)** Graphs depict AlphaFold2 pLDDT scores across residue positions for each SR-B. AlphaFold2 via ColabFold generates five protein structure models per input protein sequence and ranks each by global accuracy using the pTM metric (predicted Template-Modeling score). Line color indicates the model rank, with rank 1 being the best.

**Figure 5:**
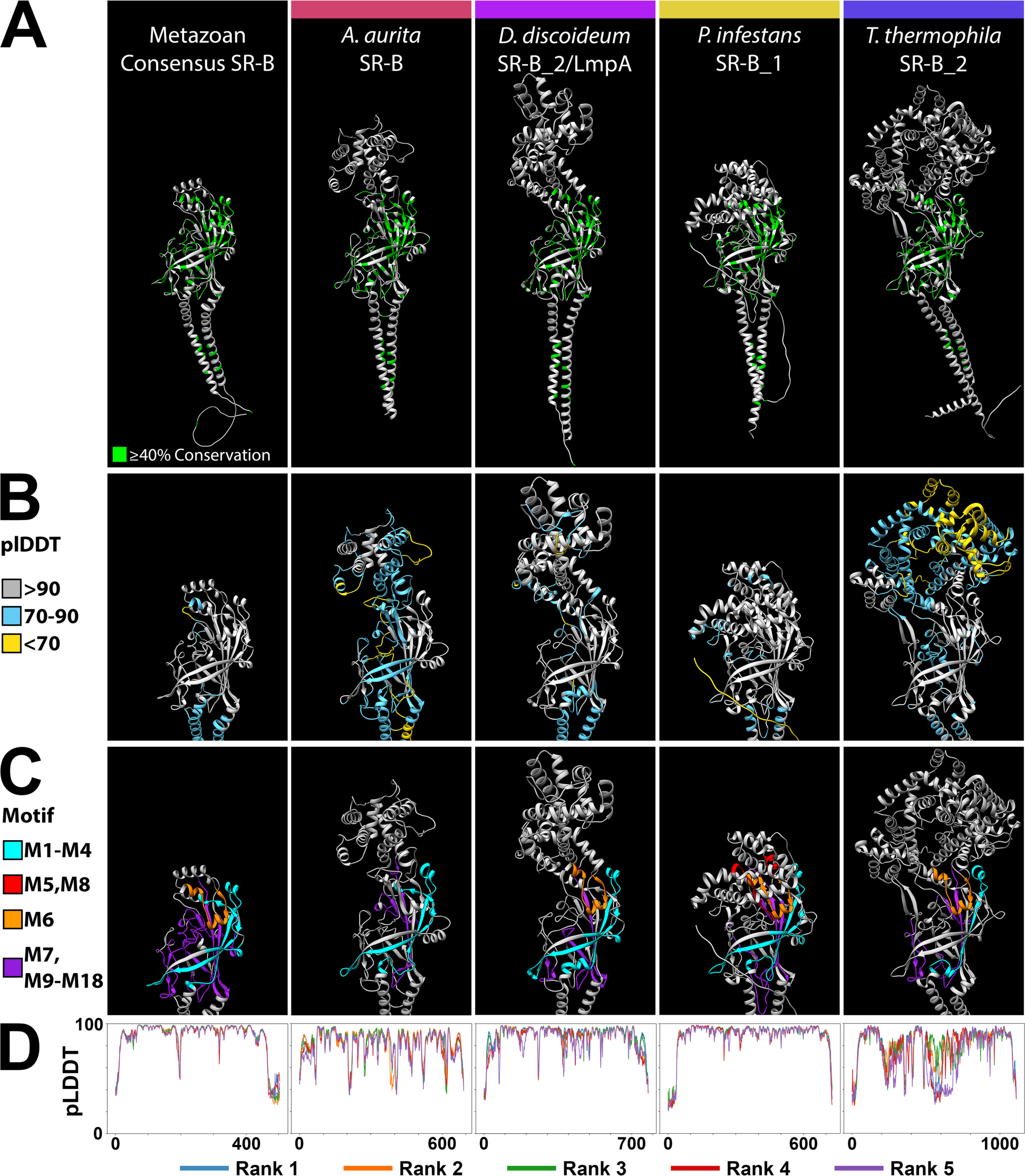
Divergent eukaryotic SR-Bs retain a conserved CD36 domain beta barrel core with lineage-specific expansions in the apex region. The highest ranked AlphaFold2 structure predictions for four exemplar eukaryote SR-B proteins with divergent, lineage-specific alpha helix structures at the apex of the CD36 ectodomain are shown, from left to right, the metazoan consensus structure, *A. aurita* SR-B, *D. discoideum* SR-B_2 (LmpA), *P. infestans* SR-B_1, and *T. thermophila* SR-B_2. Protein structure visualization was in Chimera. **A)** Green residues reflect ≥40% conservation across the alignment of all eukaryotic SR-Bs. **B)** pLDDT scores (predicted Local Distance Difference Test) computed by AlphaFold2 assess per-residue confidence across the CD36 ectodomain region of each model. Residue color reflects high confidence (gray), moderate confidence (light blue), and low confidence (yellow) pLDDT (Varadi et al., 2021). **C)** The CD36 ectodomain region of each model is annotated with the amino acid motifs discovered via STREME. Motifs M1–M4 (cyan) lie more N-terminal to the three-helix bundle at the apex, and motifs M7, M9–M18 (purple) lie more C-terminal to the three-helix bundle at the apex. Motif M6 (orange) connects two apex α-helices but is not part of the bundle itself. M5 and M8 represent the two lowest significance STREME motifs and are short motifs found within the lineage-specific sequence expansions at the apex of the CD36 ectodomain. **D)** Graphs depict AlphaFold2 pLDDT scores across residue positions for each SR-B. AlphaFold2 via ColabFold generates five protein structure models per input protein sequence and ranks each by global accuracy using the pTM metric (predicted Template-Modeling score). Line color indicates the model rank, with rank 1 being the best.

Several motifs with restricted distribution patterns across eukaryotes were also recovered. For example, motif M7 is particularly prevalent among bilaterian SR-Bs (Figure 3B) and characterizes a flexible loop region connecting the three-helix bundle at the apex to the asymmetric beta barrel of the CD36 ectodomain (Supplementary Figure 5). The two lowest scoring motifs, M5 and M8, exhibit a primarily non-metazoan distribution, with high representation among the oomycete stramenopiles.

### A structurally homologous asymmetric beta barrel is characteristic of eukaryotic SR-Bs

Divergent primary amino acid sequences can produce highly similar structures (Yang and Honig 2000; Illergård et al., 2009). Our observation of low primary sequence identity across CD36 ectodomains of eukaryotic SR-Bs (∼13%; Supplementary File 3) prompted us to interrogate SR-B protein structure. We generated AlphaFold2 (Jumper et al., 2021) and RoseTTAFold (Baek et al., 2021) protein structure models for the complete set of 279 SR-Bs (Supplementary File 9; Supplementary File 10). The relative accuracy of SR-B structure predictions was assessed by comparing the *Homo sapiens* SCARB3 X-ray crystallographic model (PDB 5LGD) against AlphaFold2 and RoseTTAFold models (Hsieh et al., 2016; Supplementary Figure 1B). Our AlphaFold2 analysis accurately predicted SCARB3 secondary and tertiary structural features (RMSD = 0.515Å across all atom pairs). RoseTTAFold also predicts most SCARB3 secondary and tertiary features, though with lower positional accuracy in the loop and alpha-helix regions when compared to the AlphaFold2 model (RMSD = 1.298Å across all atom pairs; Supplementary Figure 1C).

The relative length consistency of metazoan CD36 ectodomains also prompted us to derive a consensus sequence from an alignment of metazoan SR-Bs (Supplementary File 7; Supplementary File 8) to investigate structural similarities across metazoan SR-Bs (Supplementary Figure 6A). The consensus metazoan CD36 ectodomain is 401 aa in length and possesses alpha helices structurally congruent with the three helix bundle at the apex of bilaterian CD36 ectodomains (Supplementary Figure 6B-F). We leveraged the metazoan consensus SR-B structural model for comparative purposes.

Despite significant primary sequence variation across eukaryotic SR-Bs, all protein structure models recovered an asymmetric beta barrel, initially described in bilaterians, as the most highly conserved tertiary structure (Neculai et al., 2013; Gomez-Diaz et al., 2016; Figure 4; Figure 5). In particular, the arrangement of antiparallel β-strands is spatially consistent. The majority of conserved sequence motifs are located within the canonical asymmetric beta barrel. For example, sequence motifs M1, M2, M17 and M18 map to the longest β-strands that are also spatially adjacent to one another (Supplementary Figure 5; Figure 4C; Figure 5C). Strikingly, across eukaryotes, we also recover evidence for a conserved pocket, cavity and intramolecular tunnel feature associated with the canonical asymmetric beta barrel similar to mammalian and insect SR-Bs (Supplementary Figure 7).

To investigate putative ligand-receptor interactions with structural features of the CD36 asymmetric beta barrel, we queried SR-B AlphaFold2 models with the AlphaFill algorithm (Hekkelman et al., 2023). The AlphaFill databank contains several thousand small molecule compounds, including two known ligands for human SCARB2: cholesterol (CLR) and phosphatidylcholine (PC/PCW) (Conrad et al., 2017; Heybrock et al., 2019). CLR was recovered as a putative ligand within the intramolecular tunnel associated with the conserved asymmetric beta barrel and PCW was recovered as putative ligand of the binding pocket adjacent to the conserved asymmetric beta barrel for 90% of metazoan SR-B structural models, including the metazoan consensus model (Supplementary Table 8). Among non-metazoan SR-B structural models, CLR and PCW were recovered as putative ligands for 58% and 46% of orthologs, respectively (Supplementary Table 8). Notably, all major eukaryotic clades possess at least one recovered SR-B that recognizes CLR as a putative ligand within the intramolecular tunnel associated with the structurally conserved asymmetric beta barrel (Supplementary Table 8).

### The apex region of the CD36 domain is structurally variable across eukaryotic SR-Bs

Our analyses uncovered appreciable length-dependent structural variation across eukaryotic lineages associated with the CD36 apex (Figure 1C). Notably, eukaryotic SR-B homologs with CD36 ectodomains >500 aa have lineage-specific sequence expansions with structurally diverse apex regions (Figure 1C; Figure 5; Supplementary File 9; Supplementary File 10). AlphaFold2 and RoseTTAFold modeling of these divergent apex structures support a structural composition of multiple α-helices, with relatively low spatial confidence in isolated cases (Figure 5B). In all cases, lineage-specific sequence expansions occur within the same relative position between structural features associated with motifs M3 and M6, and correspond spatially with helices α4 and α5 of the ligand-interacting three α-helix bundle in human SCARB3 (Supplementary Figure 1A). For example, motifs M4 and M5 correspond with novel apex region α-helical structure where detected (Figure 3; Figure 5C; Figure 6). In some lineages, such as the peronosporalean oomycetes, sequence expansions are also present proximal to motif M6, spatially corresponding with α7, the third helix of the three helix bundle in human SCARB3 (Figure 6C; Supplementary Figure 1A). Notably, among divergent SR-B amino acid sequences lacking motif M6, such as the sole SR-B from the cnidarian *Aurelia aurita*, tertiary structure with high spatial congruence to motif M6 was still observed (Supplementary Figure 5; Figure 5C).

**Figure 6.**
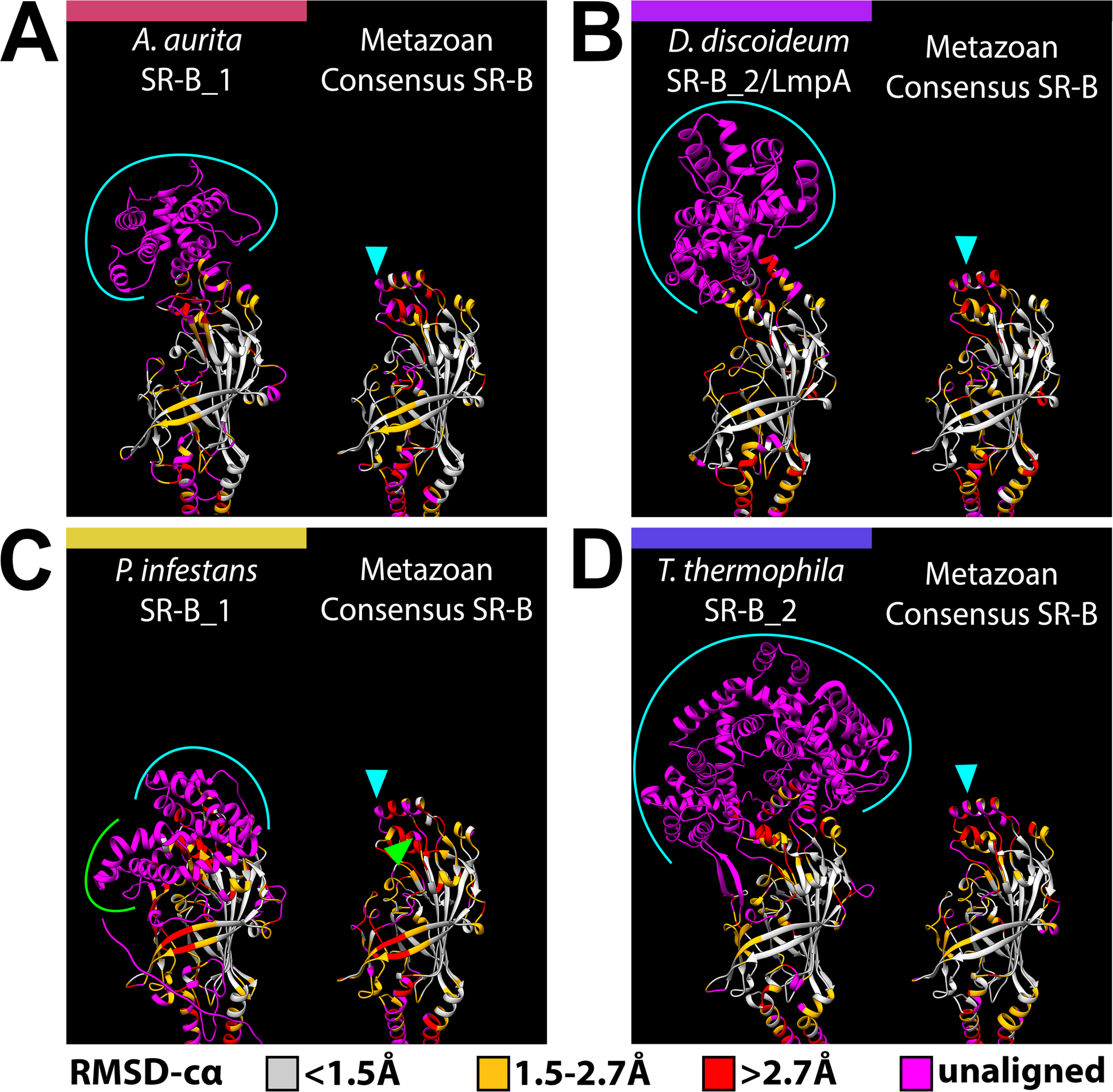
Sequence expansions coincident with the SR-B apex are divergent and have independent origins. AlphaFold2 structure predictions for four eukaryotic SR-B exemplars with lineage-specific apex structures, **A)** *Aurelia aurita* SR-B 1, **B)** *Dictyostelium discoideum* SR-B_2 (LmpA), **C)** *Phytophthora infestans* SR-B_1, and **D)** *Tetrahymena thermophila* SR-B_2 are each presented alongside the metazoan consensus SR-B structure, which serves as a structural proxy for the ‘three helix bundle’ SR-B apex. Structural superpositions were first performed for each exemplar SR-B with the metazoan consensus SR-B sequence, and alignments were generated based on the most spatially adjacent residues for each superposition. Residues in each model are color coded by the RMSD-cα (root mean square deviation between the alpha-carbons in each residue pair) to visualize structurally conserved vs divergent regions. RMSD-cα values are binned according to the categories of “high” (<1.5Å), “medium” (1.5-2.7Å), and “low” (>2.7Å) resolution used to assess the quality of crystallographic protein models (Wlodawer et al., 2007). Residues that fail to align are “unaligned” and colored magenta. Brackets highlight stretches of unaligned, lineage-specific residues that correspond with arrowheads in the metazoan consensus SR-B and represent the location of sequence expansion. Positionally, lineage-specific apex sequence expansions are consistently found to correspond with the three-helix bundle at the apex of the canonical SR-B structure.

Among SR-Bs with CD36 ectodomains <500 aa, the apex region is characterized by similar tertiary structure despite relatively low amino acid sequence conservation. None of the identified amino acid sequence motifs contribute to the three-helix bundle in eukaryotic CD36 ectodomains <500 aa. The hinge region that appears to coordinate the three helix bundle is structurally composed of a short beta-strand and helix (α6) corresponding with motif M6 and is highly conserved (Figure 3B; Supplementary Figure 5). Notably, in all cases, eukaryotic CD36 ectodomains <500 aa have apex region α-helices that correspond positionally with α4, α5, and α7 of human SCARB3 (Supplementary Figure 1A; Supplementary Figure 8). Structural comparisons reveal some variation in α-helix length and orientation. For example, in the stramenopile *Cafeteria roenbergensis* SR-B_1, two helices appear positionally in place of helix α5 (Supplementary Figure 8B). Thus our structural analyses also highlight instances of significant primary sequence divergence despite tertiary structure conservation associated with presumptive ligand interacting features of the CD36 ectodomain.

## Discussion

### Evidence for a common eukaryotic SR-B origin, with independent losses

The SR-B proteins are broadly represented across diverse eukaryotic lineages, including in some of the earliest-diverging branches (Discoba, Hemimastigophora, Ancyromonadida, CRuMs), supporting an ancient origin for the SR-B family (Figure 1B). Among some eukaryotic clades, derived taxa appear to lack CD36 domain signatures in their proteomes. For example, within the opisthokont groups Ichthyosporea and Holomycota, only early-diverging taxa retain SR-Bs. Thus we infer independent loss of SR-B homologs during the early diversification of the ichthyosporeans and in holomycotans following the divergence of the nucleariids (e.g. *Fonticula alba* and *Parvularia atlantis*). We detected similar losses within Archaeplastida, where SR-B homologs can be found within chlorophytes but not among representative streptophyte, glaucophyte, or rhodophyte species. Of note, metazoan sequences remain overrepresented in our phylogenetic analyses, highlighting the need for continued expansion of pan-eukaryotic taxon sampling and genomic resource development.

Structural congruence across eukaryotic SR-B homologs is consistent with the established position that protein tertiary structure typically demonstrates higher conservation than primary amino acid sequence identity (Yang and Honig, 2000; Illergard et al., 2009; Figure 4; Figure 5). In particular, the CD36 domain retains an asymmetric beta barrel, an adjacent binding pocket, and a connected intramolecular tunnel that are broadly shared structural features across all eukaryotic SR-B proteins (Supplementary Figure 7). The longest beta strands of the canonical asymmetric beta barrel correspond with sequence motifs that are widely distributed across eukaryotes (M1, M2, M17), further supporting an ancient origin. Additionally, ligand recognition modeling suggests cholesterol interaction with the intramolecular tunnel may represent a highly conserved feature of the SR-B receptor family across eukaryotic lineages.

### Evidence for lineage-specific CD36 ectodomain sequence expansions reflecting apex region tertiary structure diversity

The membrane-distal apex of the CD36 ectodomain is associated with ligand recognition by a three α-helix bundle and its functional role has been investigated most thoroughly in mammalian SR-B homologs (Kar et al., 2008; Kuda et al., 2013; Neculai et al., 2013; Hsieh et al., 2016). Our protein models identified structurally homologous three α-helix bundles at the apex of both metazoan and non-metazoan SR-B homologs that are <500 aa in length (Figure 1C; Supplementary File 9; Supplementary File 10). For example, similar three α-helix bundles are present in SR-B homologs found in the early branching discoban and hemimastigophran lineages (Supplementary Figure 8). We hypothesize that the ancestral CD36 ectodomain apex present in LECA may have resembled the three α-helix bundle structure present in most metazoan SR-B homologs.

Strikingly, structural comparisons between CD36 ectodomains reveal a consistent ‘deviation’ from the putative ancestral three α-helix bundle (exemplified by the metazoan consensus SR-B) positionally corresponding to α4 and α5 (Figure 6). We identified several lineage-specific sequence expansions at the membrane-distal apices of SR-B proteins >500 aa in length composed of multiple α-helices that appear to lack homology to known protein domains. We posit that these CD36 apex sequence expansions have occurred independently multiple times across Eukarya. Critically, the α5 helix motif contains amino acid residues responsible for initial ligand recognition, such as binding of long chain fatty acids (LCFAs), oxidized low-density lipoprotein (oxLDL), and *Plasmodium falciparum* MAMPs by human SCARB3 (Neculai et al., 2013; Hsieh et al., 2016). We hypothesize that a combination of lineage-specific sequence expansion and structural variation observed in the apex regions of divergent SR-B homologs may reflect the evolution of lineage-specific SR-B ligand sensing specificity.

## Conclusion

Sensing and processing environmental signals is central for many biological processes, including nutrient uptake, reproduction, and innate immunity. Integral receptor proteins at the interface of the cell and external environment are largely responsible for recognizing proximal extracellular signals and initiating cellular responses, such as signal transduction and/or interactions with co-receptors. Our protein structure data suggest that environmental and physiological ligand interactions may have driven structural variation in the membrane-distal apex region of the CD36 domain across the SR-B receptor family. The formation of alternative tertiary protein structures at the CD36 ectodomain apex may be, in part, constrained by sequence length. Lineage-specific sequence expansions in the putative ligand sensing apex appear to have contributed to the generation of novel tertiary structures. We hypothesize that these novel, lineage-specific, apex tertiary structures may correlate with ligand interactions and provide a broader understanding of the evolution of this ancient receptor family across Eukarya. Our incorporation of protein structure modeling highlights the utility of using comparative protein structure analyses to derive evolutionary inferences beyond comparison of primary amino acid sequences.

## Materials and Methods

### SR-B Identification Pipeline

The HMMER v3.3.2 package was used to search 165 publicly available genomes and transcriptomes from a diverse range of eukaryotic lineages (Supplementary Table 1) (Eddy, 1998). The hidden Markov model (*hmm*) for the Pfam CD36 domain, PF01130, was used to query predicted protein sequences from each taxon with the *hmmsearch*command (Mistry et al., 2020). Sequences that exceeded the default inclusion threshold for *hmmsearch* were retained. The CD-HIT v4.8.1 package was then used to remove identical protein sequences within each taxon (Fu et al., 2012). For each taxon, a combination of MAFFT v7.450 multiple sequence alignments (MSAs) (Katoh & Stanley, 2013), FastTree v2.1.12 (Price et al., 2010), and genome browsers (Supplementary Table 1) were used to identify likely sequence isoforms. Where multiple isoforms were detected, only the most complete representative sequence was retained for subsequent analyses.

TMHMM v2.0 (Krogh et al., 2001) and InterProScan v2.0 (Quevillon et al., 2005) were then used to identify transmembrane helices flanking the CD36 domain in order to confirm SR-B-like architecture (TM-CD36-TM). Only sequences with identifiable SR-B-like architectures were used for subsequent alignments and analyses (Supplementary File 1). Where applicable, we corroborated computationally predicted gene models that appeared to be gene model fusions or incomplete proteins with expression data in the form of transcripts or transcriptome data (e.g. Richter et al., 2018; Moreland et al., 2020). See Supplementary Table 2 for search results which specify modified sequences as well as all unique CD36 domain-containing proteins from each taxon, including those without an SR-B-like architecture.

### CD36 Domain Phylogenetic Analyses

Full length CD36 ectodomains were isolated from SR-B sequences by trimming predicted transmembrane regions and cytoplasmic tails (Supplementary File 2). For protein sequences with multiple CD36 ectodomains, only the first CD36 ectodomain was used in subsequent alignments (Trica_SR-B_1, Xensp_SR-B_1, and Pytin_SR-B_3). Recovered CD36 domains were then aligned with MAFFT v7.450 (Katoh & Standley, 2013) using the L-INS-i algorithm and the BLOSUM45 scoring matrix, with gap open penalty of 1.53 (default) and offset value of 0.123 (default) in Geneious Prime 2023.1.2 (Supplementary File 3). Identical settings were used to generate the metazoan-only MAFFT alignment (Supplementary File 6) and the cnidarian-lophotrochozoan MAFFT alignment (Supplementary Figure 3). The alignment filtering tool, trimAl v1.2 (Capella-Gutiérrez et al., 2009), was used to remove blocks of poorly aligned residues in the CD36 ectodomain alignment to generate a filtered alignment optimized for phylogenetic signal using the “-gappyout” option (Supplementary File 4). ModelTest-NG v0.1.7 was used to identify LG+G4+F as the best-fit amino acid substitution model for the filtered alignment (Darriba et al., 2019). Maximum likelihood (ML) analyses were performed using the Pthreads version of RAxML v8.2.12 (Stamatakis, 2014). A total of 300 independent ML searches were performed on randomized maximum parsimony starting trees. 100 bootstrap replicates were performed and used to draw bipartitions on the best scoring tree (log-likelihood score of -223409.574461) (Supplementary Figure 2 and Supplementary File 5).

Bayesian analyses were performed with MrBayes v3.2.7 (Ronquist and Huelsenbeck, 2003). Initially two independent runs of 5 million generations with five chains each using default heating and the “LG+G4” amino acid model were performed. The average standard deviation of split frequencies between the initial two runs was 0.147471 reflecting a lack of convergence. Therefore, an additional 1 million runs were performed, for a total of 6 million generations, to assess the feasibility of achieving convergence between the two runs. The average standard deviation of split frequencies between the two 6 million runs was 0.145354, thus the Bayesian analyses failed to converge. To attempt to recover supported nodes despite the lack of convergence, the initial 25% of trees from each run were removed as burn-in and posterior probabilities (BPP) were calculated (Supplementary Figure 3 and Supplementary File 6). FigTree v1.4.4 (http://tree.bio.ed.ac.uk/software/figtree/) was used to visualize both ML and Bayesian trees.

### SR-B Motif Prediction

Amino acid motifs were identified using STREME v5.5.3 (Bailey, 2021). Default motif length settings were used (minimum and maximum motif length of 8 and 30 amino acids respectively). The STREME algorithm stops searching for motifs after three calls when the default p-value threshold exceeds 0.05. Recovered motifs are evaluated for statistical significance by STREME by determining the probability of discriminating primary sequences from a subset of randomized and shuffled control sequences. When STREME reports more than one motif, the p-value does not completely account for multiple testing, in this case E-values are calculated for each motif to evaluate statistical significance (Bailey, 2021). The strength of a putative motif match in a given sequence is determined by deriving a ‘sequence score’ by summing the weight of each position in the motif PWM (Bailey, 2021). STREME computes a match threshold for each motif, sequence elements with scores below the match threshold are disregarded.

### SR-B Structure Prediction

Complete SR-B sequences were modeled with AlphaFold2 (Jumper et al., 2021) via the ColabFold web server (accessed 3/31/22–12/15/22 ; Mirdita et al., 2022) and with RoseTTAFold (Baek et al., 2021) via the Robetta web server (accessed 3/14/22–12/15/22; Kim et al., 2004). Sequences greater than 1401 amino acids were excluded from both AlphaFold2 and RoseTTAFold structure prediction (ColabFold and Robetta web server parameters require sequences less than 1401 amino acids in length). Protein sequences containing undetermined amino acid positions (X) were replaced with alanine (A), a non-bulky, non-reactive amino acid residue commonly used in protein structure and stability experiments (Matthews, 1996). AlphaFold2 via ColabFold generated 5 models for each SR-B sequence, ranking each by global accuracy with a predicted template-modeling score (pTM) (Jumper et al., 2021), an approximation of the “true” template modeling score (TM-score) (Zhang and Skolnick, 2004). pTM scores range from 0 to 1.0, with 1.0 being the best. Models with a pTM of >0.7 were used for comparative analyses. The highest ranked model above this pTM threshold for each SR-B was used for downstream analyses.

Local accuracy was measured with the predicted Local Distance Difference Test (pLDDT) computed by AlphaFold which gives per-residue confidence scores (Supplementary Table 5). pLDDT values range from 0–100, with 100 being the best. SR-B RoseTTAFold models were evaluated with the local distance difference test (lDDT) using the deep learning framework DeepAccNet (Mariani et al., 2013; Hiranuma et al., 2021). lDDT scores range from 0 to 1.0, with 1.0 being the best. Local accuracy was assessed by root mean square (RMS) error of α-carbons in Ångstroms (Supplementary Table 5). Internal cavities were predicted with the CASTp 3.0 web server (Tian et al., 2018) using a probe radius of 0.9Å. All protein models were visualized in UCSF Chimera v1.17 (Petterson et al., 2004).

An additional alignment of complete metazoan SR-B sequences was generated to produce a metazoan consensus SR-B sequence (Supplementary File 6; Supplementary File 7). The metazoan consensus SR-B sequence was fed into both AlphaFold2 and RoseTTAFold structure prediction pipelines to yield a metazoan consensus protein structure prediction (Supplementary Figure 6).

## Supporting information

Spplementary Figures and Tables

## Acknowledgments

This work was supported by the National Science Foundation [grant number 2013692].

## Author Contributions

Conceptualization, W.E.B.; Methodology, R.T.B. and W.E.B.; Formal Analysis and Investigation, R.T.B. and W.E.B.; Data Curation, R.T.B. and W.E.B; Visualization, R.T.B. and W.E.B.; Writing - Original Draft, R.T.B., L.E.V. and W.E.B.; Writing - Review and Editing, All authors; Resources, W.E.B.; Supervision, W.E.B.; Project Administration, W.E.B.; Funding Acquisition, N.T.-K. and W.E.B.

## Data Availability

The data underlying this article are available in the article and in its online supplementary material.

## Notes

### Competing Interest Statement

The authors have declared no competing interest.

## References

Ausubel FM. Are innate immune signaling pathways in plants and animals conserved? Nature Immunology 2005; 6(10):973–979. 10.1038/ni1253.

Baek M, DiMaio F, Anishchenko I, Dauparas J, Ovchinnikov S, Lee Gyu R, Wang J, Cong Q, Kinch Lisa N, Schaeffer RD, Millan C, Park H, Adams C, Glassman CR, Degiovanni A, Pereira JH, Rodrigues AV, van Dijk AA, Ebrecht AC, Opperman DJ, Sagmeister T, Buhlheller C, Pavkov-Keller T, Rathinaswamy MK, Dalwadi U, Yip CK, Burke JE, Garcia KC, Grishin NV, Adams PD, Read RJ, Baker D. Accurate prediction of protein structures and interactions using a three-track neural network. Science 2021; 373(6557):871–876.10.1126/science.abj8754.

Bailey T. 2021. STREME: accurate and versatile sequence motif discovery. Bioinformatics 37(18):2834–2840. 10.1093/bioinformatics/btab203

Bi W-J, Li D-X, Xu Y-H, Xu S, Li J, Zhao X-F, Wang J-X. 2015. Scavenger receptor B protects shrimp from bacteria by enhancing phagocytosis and regulating expression of antimicrobial peptides. Developmental & Comparative Immunology 51(1):10–21. 10.1016/j.dci.2015.02.001.

Burki F, Roger AJ, Brown MW, Simpson AGB. 2020. The New Tree of Eukaryotes. Trends in Ecology & Evolution 35(1):43–55. 10.1016/j.tree.2019.08.008.

Capella-Gutiérrez S, Silla-Martínez JM, Gabaldón T. 2009. trimAl: a tool for automated alignment trimming in large-scale phylogenetic analyses. Bioinformatics 25(15):1972–1973. 10.1093/bioinformatics/btp348.

Carr M, Richter DJ, Fozouni P, Smith TJ, Jeuck A, Leadbeater BSC, Nitsche F. 2017. A six-gene phylogeny provides new insights into choanoflagellate evolution. Molecular Phylogenetics and Evolution 107:166–178.10.1016/j.ympev.2016.10.011.

Chen Y, Zhang J, Cui W, Silverstein RL. 2022. CD36, a signaling receptor and fatty acid transporter that regulates immune cell metabolism and fate. J Exp Med 219(6):e20211314. 10.1084/jem.20211314

Chovancova E, Pavelka A, Benes P, Strnad O, Brezovsky J, Kozlikova B, Gora A, Sustr V, Klvana M, Medek P, Biedermannova L, Socho J, Damborsky J. 2012. Caver 3.0: A Tool for the Analysis of Transport Pathways in Dynamic Protein Structures. PLOS Computational Biology 8(10): e1002708. 10.1371/journal.pcbi.1002708

Conrad KS, Cheng T-W, Ysselstein D, Heybrock S, Hoth LR, Chrunyk BA, am Ende CW, Krainc D, Scwake M, Saftig P, Liu S, Qui X, Ehlers MD. Lysosomal integral membrane protein-2 as a phospholipid receptor revealed by biophysical and cellular studies. Nature Communications 2017; 8:1908. 10.1038/s41467-017-02044-8

Darriba D, Posada D, Kozlov AM, Stamatakis A, Morel B, Flouri T. 2020. ModelTest-NG: A New and Scalable Tool for the Selection of DNA and Protein Evolutionary Models. Molecular Biology and Evolution 37(1):291–294. 10.1093/molbev/msz189.

Derelle R, López-García P, Timpano H, Moreira D. 2016. A Phylogenomic Framework to Study the Diversity and Evolution of Stramenopiles (=Heterokonts). Molecular Biology and Evolution 33(11):2890–2898. 10.1093/molbev/msw168.

Dinguirard N, Yoshino TP. 2006. Potential role of a CD36-like class B scavenger receptor in the binding of modified low-density lipoprotein (acLDL) to the tegumental surface of Schistosoma mansoni sporocysts. Molecular and Biochemical Parasitology 146(2):219–230. 10.1016/j.molbiopara.2005.12.010.

Dobri, A.M., Dudău, M., Enciu, A.M. and Hinescu, M.E., 2021. CD36 in Alzheimer’s disease: an overview of molecular mechanisms and therapeutic targeting. Neuroscience, 453, pp.301–311. 10.1016/j.neuroscience.2020.11.003

Eddy SR. 1998. Profile hidden Markov models. Bioinformatics 14(9):755–763.10.1093/bioinformatics/14.9.755.

Erdman LK, Cosio G, Helmers AJ, Gowda DC, Grinstein S, Kain KC. 2009. CD36 and TLR Interactions in Inflammation and Phagocytosis: Implications for Malaria. J Immunol 183(10):6452–6459. 10.4049/jimmunol.0901374

Ettahi K, Lhee D, Sung JY, Simpson AGB, Park JS, Yoon HS. 2021. Evolutionary History of Mitochondrial Genomes in Discoba, Including the Extreme Halophile Pleurostomum flabellatum (Heterolobosea). Genome Biology and Evolution 13(2):evaa241. 10.1093/gbe/evaa241.

Franc NC, Dimarcq J-L, Lagueux M, Hoffmann J, Ezekowitz RAB. Croquemort, A Novel Drosophila Hemocyte/Macrophage Receptor that Recognizes Apoptotic Cells. Immunity 1996; 4(5):431–443.10.1016/S1074-7613(00)80410-0.

Fu L, Niu B, Zhu Z, Wu S, Li W. CD-HIT: accelerated for clustering the next-generation sequencing data. Bioinformatics 2012; 28(23):3150–3152. 10.1093/bioinformatics/bts565.

Glatz JFC, Luiken JJFP. Dynamic role of the transmembrane glycoprotein CD36 (SR-B2) in celluar fatty acid uptake and utilization. Journal of Lipid Research 2018; 59:R082933. 10.1194/jlr.R082933

Gomez-Diaz C, Bargeton B, Abuin L, Bukar N, Reina JH, Bartoi T, Graf M, Ong H, Ulbrich MH, Masson J-F, Benton R. A CD36 ectodomain mediates insect pheromone detection via a putative tunnelling mechanism. Nature Communications 2016; 7(1):11866. 10.1038/ncomms11866.

Hekkelman ML, de Vries I, Joosten RP, Perrakis A. AlphaFill: enriching AlphaFold models ligands and cofactors. Nature Methods 2023; 20: 205–213. 10.1038/s41592-022-01685-y

Heybrock S, Kanerva K, Meng Y, Ing C, Liang A, Xiong Z-J, Weng X, Kim YA, Collins R, Trimble W, Pomès R, Privé GG, Annaert W, Schwake M, Heeren J, Lüllmann-Rauch R, Grinstein S, Ikonen E, Saftig P, Neculai D. Lysosomal integral membrane protein-2 (LIMP-2/SCARB2) is involved in lysosomal cholesterol export. Nature Communications 2019; 10:3521. 10.1038/s41467-019-11425-0

Hiranuma N, Park H, Baek M, Anishchenko I, Dauparas J, Baker D. Improved protein structure refinement guided by deep learning based accuracy estimation. Nature Communications 2021;12(1):1340. 10.1038/s41467-021-21511-x

Hsieh F-L, Turner L, Bolla JR, Robinson CV, Lavstsen T, Higgins MK. The structural basis for CD36 binding by the malaria parasite. Nature Communications 2016;7(1):12837. 10.1038/ncomms12837

Illergård K, Ardell DH, Elofsson A. 2009. Structure is three to ten times more conserved than sequence—A study of structural response in protein cores. Proteins: Structure, Function and Bioinformatics 77(3): 499–508. 10.1002/prot.22458

Jay AG, Chen AN, Paz MA, Hung JP, Hamilton JA. CD36 Binds Oxidized Low Density Lipoprotein (LDL) in a mechanism Dependent upon Fatty Acid Binding. Journal of Biological Chemistry 290(8):4590–4603. 10.1074/jbc.M114.627026

Jumper J, Evans R, Pritzel A, Green T, Figurnov M, Ronneberger O, Tunyasuvunakool K, Bates R, Žídek A, Potapenko A, Bridgland A, Meyer C, Kohl SAA, Ballard AJ, Cowie A, Romera-Paredes B, Nikolov S, Jain R, Adler J, Back T, Petersen S, Reiman D, Clancy E, Zielinski M, Steinegger M, Pacholska M, Berghammer T, Bodenstein S, Silver D, Vinyals O, Senior AW, Kavukcuoglu K, Kohli P, Hassabis D. 2021. Highly accurate protein structure prediction with AlphaFold. Nature 596(7873):583–589. 10.1038/s41586-021-03819-2.

Kar NS, Ashraf MZ, Valiyaveettil M, Podrez EA. 2008. Mapping and Characterization of the Binding Site for Specific Oxidized Phospholipids and Oxidized Low Density Lipoprotein of Scavenger Receptor CD36. Journal of Biological Chemistry 283(13):8765–8771. 10.1074/jbc.M709195200

Karakesisoglou I, Janssen K-P, Eichinger L, Noegel AA, Schleicher M. 1999. Identification of a Suppressor of the Dictyostelium Profilin-minus Phenotype as a CD36/LIMP-II Homologue. Journal of Cell Biology 145(1):167–181. 10.1083/jcb.145.1.167.

Katoh K, Standley DM. 2013. MAFFT Multiple Sequence Alignment Software Version 7: Improvements in Performance and Usability. Molecular Biology and Evolution 30(4):772–780.10.1093/molbev/mst010.

Kim DE, Chivian D, Baker D. 2004. Protein structure prediction and analysis using the Robetta server. Nucleic acids research 32(Web Server issue):W526–W531. 10.1093/nar/gkh468.

Koropatnick TA, Engle JT, Apicella MA, Stabb EV, Goldman WE, McFall-Ngai MJ. 2004. Microbial Factor-Mediated Development in a Host-Bacterial Mutualism. Science 306(5699):1186–1188.10.1126/science.1102218.

Krogh A, Larsson B, von Heijne G, Sonnhammer ELL. 2001. Predicting transmembrane protein topology with a hidden markov model: application to complete genomes11Edited by F. Cohen. Journal of Molecular Biology 305(3):567–580. 10.1006/jmbi.2000.4315.

Kuda O, Pietka TA, Demianova Z, Kudova E, Cvacka J, Kopecky J. 2013. Sulfo-*N-*succinimidyl Oleate (SSO) Inhibits Fatty Acid Uptake and Signaling for Intracellular Calcium via Binding CD36 Lysine 164. Journal of Biological Chemistry 228(22):15547–15555. 10.1074/jbc.M113.473298

Li X, Hou Z, Xu C, Shi X, Yang L, Lewis LA, Zhong B. 2021. Large Phylogenomic Data sets Reveal Deep Relationships and Trait Evolution in Chlorophyte Green Algae. Genome Biology and Evolution 13(7):evab101.10.1093/gbe/evab101.

Li Y, Shen X-X, Evans B, Dunn CW, Rokas A. 2021. Rooting the Animal Tree of Life. Molecular Biology and Evolution 38(10):4322–4333. 10.1093/molbev/msab170.

Mariani V, Biasini M, Barbato A, Schwede T. 2013. lDDT: a local superposition-free score for comparing protein structures and models using distance difference tests. Bioinformatics 29(21):2722–2728. 10.1093/bioinformatics/btt473

Matthews BW. 1996. Structural and genetic analysis of the folding and function of T4 lysozyme. The FASEB Journal 10(1):35–41. 10.1096/fasebj.10.1.8566545.

Means TK, Mylonakis E, Tampakakis E, Colvin RA, Seung E, Puckett L, Tai MF, Stewart CR, Pukkila-Worley R, Hickman SE, Moore KH, Calderwood SB, Hacohen N, Luster AD, Khoury JE. 2009. Evolutionarily conserved recognition and innate immunity to fungal pathogens by the scavenger receptors SCARF1 and CD36. Journal of Experimental Medicine 206(3):637–653. 10.1084/jem.20082109.

Mirdita M, Schütze K, Moriwaki Y, Heo L, Ovchinnikov S, Steinegger M. 2022. ColabFold: making protein folding accessible to all. Nature Methods 19: 679–682. 10.1038/s41592-022-01488-1

Mistry J, Chuguransky S, Williams L, Qureshi M, Salazar Gustavo A, Sonnhammer ELL, Tosatto SCE, Paladin L, Raj S, Richardson LJ, Finn RD, Bateman A. 2021. Pfam: The protein families database in 2021. Nucleic Acids Research 49(D1):D412–D419. 10.1093/nar/gkaa913.

Moreland RT, Nguyen A, Ryan JF, Schnitzler CE, Koch BJ, Siewert K, Wolfsberg TG, Baxevanis AD. 2014. A customized Web portal for the genome of the ctenophore *Mnemiopsis leidyi*. BMC Genomics 15: 316. 10.1186/1471-2164-15-316

Moreland RT, Nguyen A, Ryan JF, Baxevanis AD. 2020. The *Mnemiopsis* Genome Project Portal: integrating new gene expression resources and improving data visualization. Database 2020: baaa029. 10.1093/database/baaa029

Müller WEG, Thakur NL, Ushijima H, Thakur AN, Krasko A, Le Pennec Gl, Indap MM, Perović-Ottstadt S, Schröder HC, Lang G, Bringmann G. 2004. Matrix-mediated canal formation in primmorphs from the sponge Suberites domuncula involves the expression of a CD36 receptor-ligand system. Journal of Cell Science 117(12):2579–2590. 10.1242/jcs.01083.

Neculai D, Schwake M, Ravichandran M, Zunke F, Collins RF, Peters J, Neculai M, Plumb J, Loppnau P, Pizarro JC, Seitova A, Trimble WS, Saftig P, Grinstein S, Dhe-Paganon S. 2013. Structure of LIMP-2 provides functional insights with implications for SR-BI and CD36. Nature 504(7478):172–176. 10.1038/nature12684.

Neubauer EF, Poole AZ, Detournay O, Weis VM, Davy SK. 2016. The scavenger receptor repertoire in six cnidarian species and its putative role in cnidarian-dinoflagellate symbiosis. PeerJ 4:e2692.10.7717/peerj.2692.

Nichols Z, Vogt RG. 2008. The SNMP/CD36 gene family in Diptera, Hymenoptera and Coleoptera: Drosophila melanogaster, D. pseudoobscura, Anopheles gambiae, Aedes aegypti, Apis mellifera, and Tribolium castaneum. Insect Biochemistry and Molecular Biology 38(4):398–415.10.1016/j.ibmb.2007.11.003.

Özbek S, Balasubramanian PG, Chiquet-Ehrismann R, Tucker RP, Adams JC. 2010. The Evolution of Extracellular Matrix. Molecular Biology of the Cell 21(24):4300–4305. 10.1091/mbc.e10-03-0251.

Papale GA, Hanson PJ, Sahoo D. 2011. Extracellular Disulfide Bonds Support Scavenger Receptor Class B Type I-Mediated Cholesterol Transport. Biochemistry 50(28):6245–6254. 10.1021/bi2005625.

Park, L., Zhou, J., Zhou, P., Pistick, R., El Jamal, S., Younkin, L., Pierce, J., Arreguin, A., Anrather, J., Younkin, S.G. and Carlson, G.A., 2013. Innate immunity receptor CD36 promotes cerebral amyloid angiopathy. Proceedings of the National Academy of Sciences, 110(8), pp.3089–3094. 10.1073/pnas.1300021110

Pettersen EF, Goddard TD, Huang CC, Couch GS, Greenblatt DM, Meng EC, Ferrin TE. 2004. UCSF Chimera—A visualization system for exploratory research and analysis. Journal of Computational Chemistry 25(13):1605–1612. 10.1002/jcc.20084.

PrabhuDas MR, Baldwin CL, Bollyky PL, Bowdish DME, Drickamer K, Febbraio M, Herz J, Kobzik L, Krieger M, Loike J, McVicker B, Means TK, Moestrup SK, Post SR, Sawamura T, Silverstein S, Speth RC, Telfer JC, Thiele GM, Wang X-Y, Wright SD, Khoury JE. 2017. A Consensus Definitive Classification of Scavenger Receptors and Their Roles in Health and Disease. The Journal of Immunology 198(10):3775. 10.4049/jimmunol.1700373.

Price MN, Dehal PS, Arkin AP. 2010. FastTree 2 – Approximately Maximum-Likelihood Trees for Large Alignments. PLOS ONE 5(3):e9490. 10.1371/journal.pone.0009490.

Quevillon E, Silventoinen V, Pillai S, Harte N, Mulder N, Apweiler R, Lopez R. 2005. InterProScan: protein domains identifier. Nucleic Acids Research 33(suppl_2):W116–W120. 10.1093/nar/gki442.

Rasmussen JT, Berglund L, Rasmussen MS, Petersen TE. Assignment of disulfide bridges in bovine CD36. European Journal of Biochemistry 1998; 257(2):488–494. 10.1046/j.1432-1327.1998.2570488.x.

Richter DJ, Fozouni P, Eisen MB, King N. 2018. Gene family innovation, conservation and loss on the animal stem lineage. eLife 7:e34226. 10.7554/eLife.34226.

Richter DJ, Levin TC. 2019. The origin and evolution of cell-intrinsic antibacterial defenses in eukaryotes. Current Opinion in Genetics & Development 58-59:111–122. 10.1016/j.gde.2019.09.002.

Richter DJ, Berney C, Strassert JFH, Yu-Ping Poh Y-P, Emily K. Herman EK, Muñoz-Gómez SA, Jeremy G.Wideman JG, Fabien Burki F, de Vargas C. EukProt: A database of genome-scale predicted proteins across the diversity of eukaryotes. Peer Community Journal 2022; 2: e56. 10.24072/pcjournal.173

Ronquist F, Huelsenbeck JP. 2003. MrBayes 3: Bayesian phylogenetic inference under mixed models. Bioinformatics 19:1572–1574. 10.1093/bioinformatics/btg180

Sattler N, Bosmani C, Barisch C, Guého A, Gopaldass N, Dias M, Leuba F, Bruckert F, Cosson P, Soldati T. 2018. Functions of the Dictyostelium LIMP-2 and CD36 homologues in bacteria uptake, phagolysosome biogenesis and host cell defense. Journal of Cell Science 131(17):jcs218040.10.1242/jcs.218040.

Schaefer L. 2014. Complexity of Danger: The Diverse Nature of Damage-associated Molecular Patterns*. Journal of Biological Chemistry 289(51):35237–35245. 10.1074/jbc.R114.619304.

Sheedy FJ, Grebe A, Rayner KJ, Kalantari P, Ramkhelawon B, Carpenter SB, Becker CE, Ediriweera HN, Mullick AE, Golenbock DT, Stuart LM, Latz E, Fitzgerald KA, Moore KJ. 2013. CD36 coordinates NLRP3 inflammasome activation by facilitating intracellular nucleation of soluble ligands into particulate ligands in sterile inflammation. Nature Immunology 14:812–820. 10.1038/ni.2639

Silverstein Roy L, Febbraio M. 2009. CD36, a Scavenger Receptor Involved in Immunity, Metabolism, Angiogenesis, and Behavior. Science Signaling 2(72):re3–re3. 10.1126/scisignal.272re3.

Stamatakis A. 2014. RAxML version 8: a tool for phylogenetic analysis and post-analysis of large phylogenies. Bioinformatics 30(9):1312–1313. 10.1093/bioinformatics/btu033.

Stewart CR, Stuart LM, Wilkinson K, van Gils JM, Deng J, Halle A, Rayner KJ, Boyer L, Zhong R, Frazier WA, Lacy-Hulbert A, Khoury JE, Golenbock DT, Moore KJ. 2010. CD36 ligands promote sterile inflammation through assembly of a Toll-like receptor 4 and 6 heterodimer. Nature Immunology 11:155–161. 10.1038/ni.1836

Strassert JFH, Irisarri I, Williams TA, Burki F. 2021. A molecular timescale for eukaryote evolution with implications for the origin of red algal-derived plastids. Nature Communications 12(1):1879.10.1038/s41467-021-22044-z

Stuart, L.M., Bell, S.A., Stewart, C.R., Silver, J.M., Richard, J., Goss, J.L., Tseng, A.A., Zhang, A., El Khoury, J.B., Moore, K.J., 2007. CD36 signals to the actin cytoskeleton and regulates microglial migration via a p130Cas complex. J. Biol. Chem. 282, 27392–27401. 10.1074/jbc.M702887200

Taban, Q., Mumtaz, P.T., Masoodi, K.Z., Haq, E. and Ahmad, S.M., 2022. Scavenger receptors in host defense: from functional aspects to mode of action. Cell Communication and Signaling, 20, pp.1–17. 10.1186/s12964-021-00812-0

Thorne RF, Meldrum CJ, Harris SJ, Dorahy DJ, Shafren DR, Berndt MC, Burns GF, Gibson PG. 1997. CD36 Forms Covalently Associated Dimers and Multimers in Platelets and Transfected COS-7 Cells. Biochemical and Biophysical Research Communications 240(3):812–818. 10.1006/bbrc.1997.7755

Tian W, Chen C, Lei X, Zhao J, Liang J. 2018. CASTp 3.0: computed atlas of surface topography of proteins. Nucleic Acids Research 46(W1):W363–367. 10.1093/nar/gky473

Varadi M, Anyango S, Deshpande M, Nair S, Natassia C, Yordanova G, Yuan D, Stroe O, Wood G, Laydon A et al. . 2021. AlphaFold Protein Structure Database: massively expanding the structural coverage of protein-sequence space with high-accuracy models. Nucleic Acids Research 50(D1):D439–D444.10.1093/nar/gkab1061.

Wheeler GL, Miranda-Saavedra D, Barton GJ. 2008. Genome Analysis of the Unicellular Green Alga Chlamydomonas reinhardtii Indicates an Ancient Evolutionary Origin for Key Pattern Recognition and Cell-Signaling Protein Families. Genetics 179(1):193–197. 10.1534/genetics.107.085936.

Wlodawer A, Minor W, Dauter Z, and Jaskolski M. 2007. Protein crystallography for non-crystallographers, or how to get the best (but not more) from published macromolecular structures. The FEBS Journal 275(1):1–21. 10.1111/j.1742-4658.2007.06178.x

Yang, A-S, Honig B. 2000. An integrated approach to the analysis and modeling of protein sequences and structures. III. A comparative study of sequence conservation in protein structural families using multiple structural alignments. Journal of Molecular Biology 301(3):691–711. 10.1006/jmbi.2000.3975

Yu M, Lau TY, Carr SA, Krieger M. 2012. Contributions of a Disulfide Bond and a Reduced Cysteine Side Chain to the Intrinsic Activity of the High-Density Lipoprotein Receptor SR-BI. Biochemistry 51(50):10044–10055. 10.1021/bi301203x.

Zhang Y, Skolnick J. 2004. Scoring function for automated assessment of protein structure template quality. Proteins: Structure, Function, and Bioinformatics 57(4):702–710. 10.1002/prot.20264

