## Supplementary material for "Phylogenetic and protein structure analyses provide insight into the evolution and diversification of the CD36 domain ‘apex’ among scavenger receptor class B proteins across Eukarya": Spplementary Figures and Tables: Supplementary Figures.pdf

### Supplementary Figures 1-8

### Supplementary Table Legends 1-7

**Supplementary Figure 1**

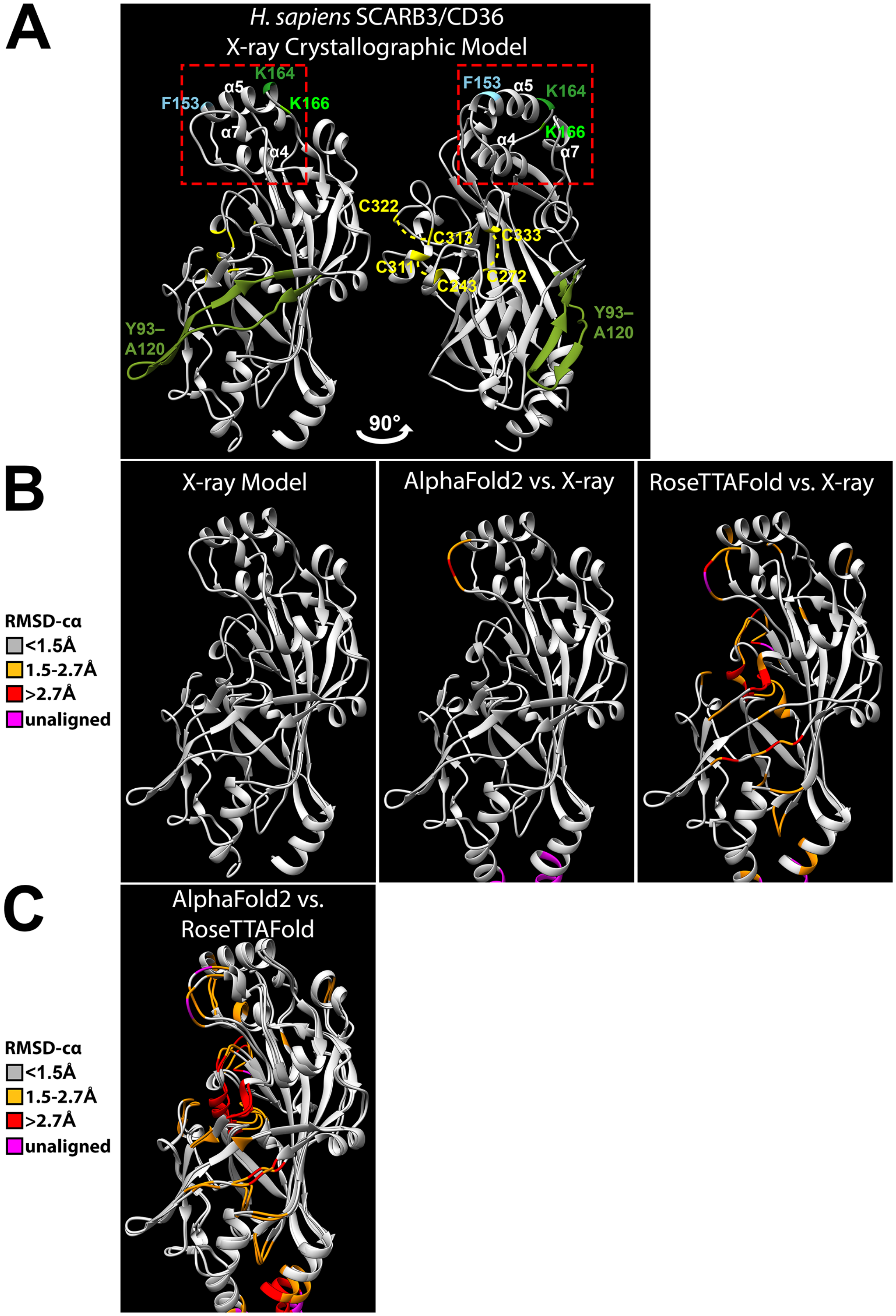

**Supplementary Figure 1: *H. sapiens* SCARB3 X-ray model annotated with significant features and validation of AlphaFold2 and RoseTTAFold models. A)** Ribbon model for *Homo sapiens* SCARB3 based on the crystal structure (Hsieh et al., 2016; PDB ID: 5LGD) annotated with residues identified in previous functional studies. Left and right are identical models rotated 90 degrees. The membrane-distal "apex" region of the protein is boxed in red, consists of a cluster of several alpha helices, and is functionally associated with ligand recognition. K164 was identified as critical for binding oxidized phosphatidylcholines specific to CD36, oxidized low-density lipoproteins (Kar et al., 2008), and fatty acids (Kuda et al., 2013). K166 was also identified as critical for binding oxidized low-density lipoproteins (Kar et al., 2008). F153 serves as the docking site for the *Plasmodium falciparum* PfEMP1 protein to recognize erythrocytes. Residues Y93–A120 correspond to the CLESH domain and bind proteins containing thrombospondin type 1 homology (TSR) domains (Silverstein et al., 2001). The three pairs of cysteines involved in disulfide bridge formation are labeled yellow on the model to the right. Disulfide bridge-1 is formed by C272–C333, disulfide bridge-2 is formed by C313–C321, and disulfide bridge-3 is formed by C242–C310. **B)** The X-ray model (left), followed by the RoseTTAFold structure prediction (middle) and the AlphaFold2 structure prediction (right) for SCARB3 structurally aligned in Chimera. Residues colored in the RoseTTAFold and AlphaFold2 structures based on the root mean square deviation of the carbon atoms (RMSD-cα) corresponding to the crystal structure. Angstrom deviation key to the left. **C)** Overlay of the RoseTTAFold and AlphaFold2 models with residues colored by RMSD-cα between corresponding atoms in each model. Angstrom deviation key to the left.

**Supplementary Figure 2**

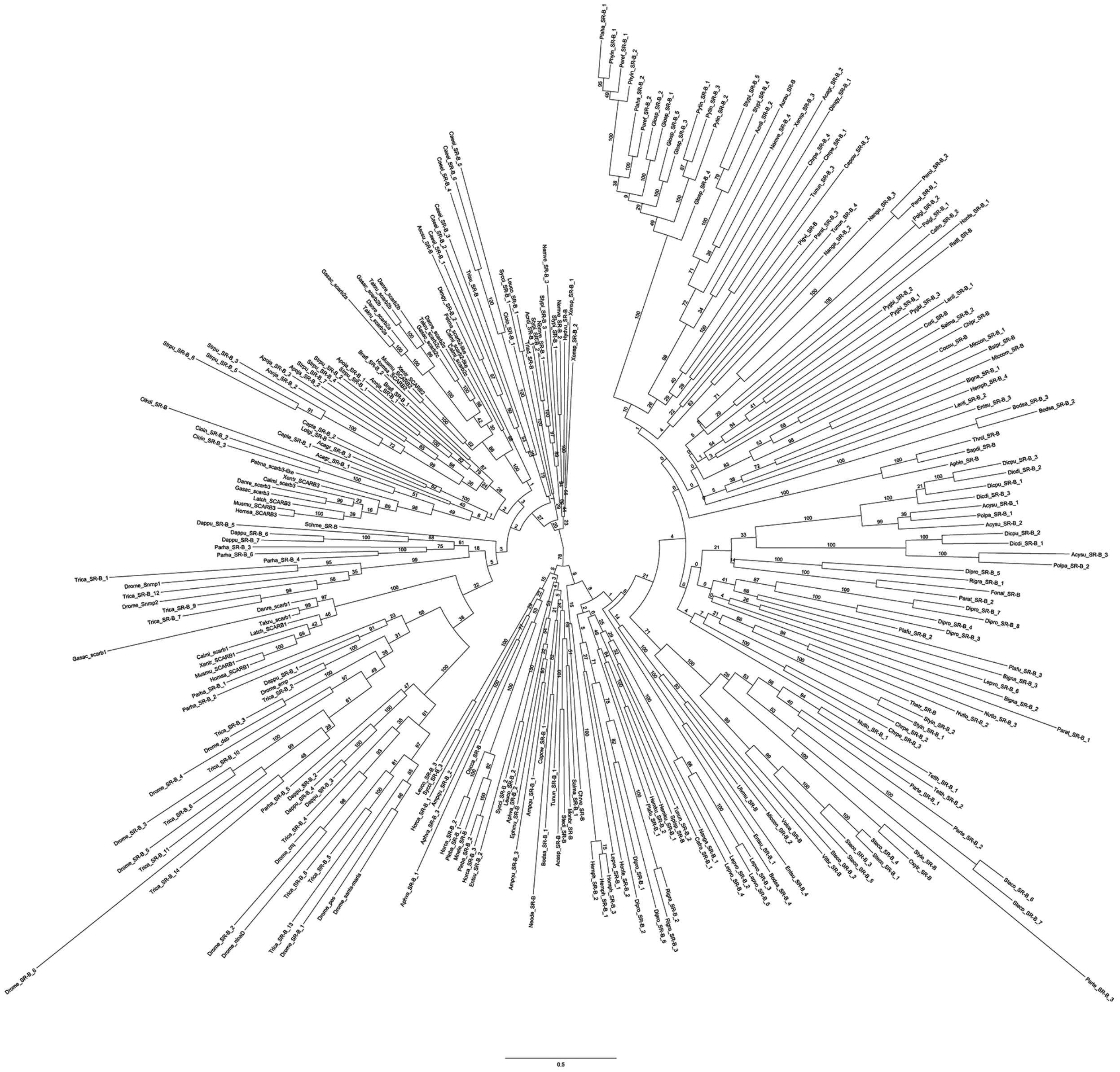

**Supplementary Figure 2: Maximum likelihood phylogenetic analysis.** Best tree based on amino acid MAFFT alignment corresponding to 279 eukaryotic CD36 ectodomains. inferred via RAxML v8.2.12 using the LG+G4+F evolutionary model and 100 bootstrap replicates. Labels indicate node bootstrap (BS) values. Tip labels are protein names. Refer to Supplementary Table 2 for source genome or transcriptome, and sequence identifiers.

### Supplementary Figure 3

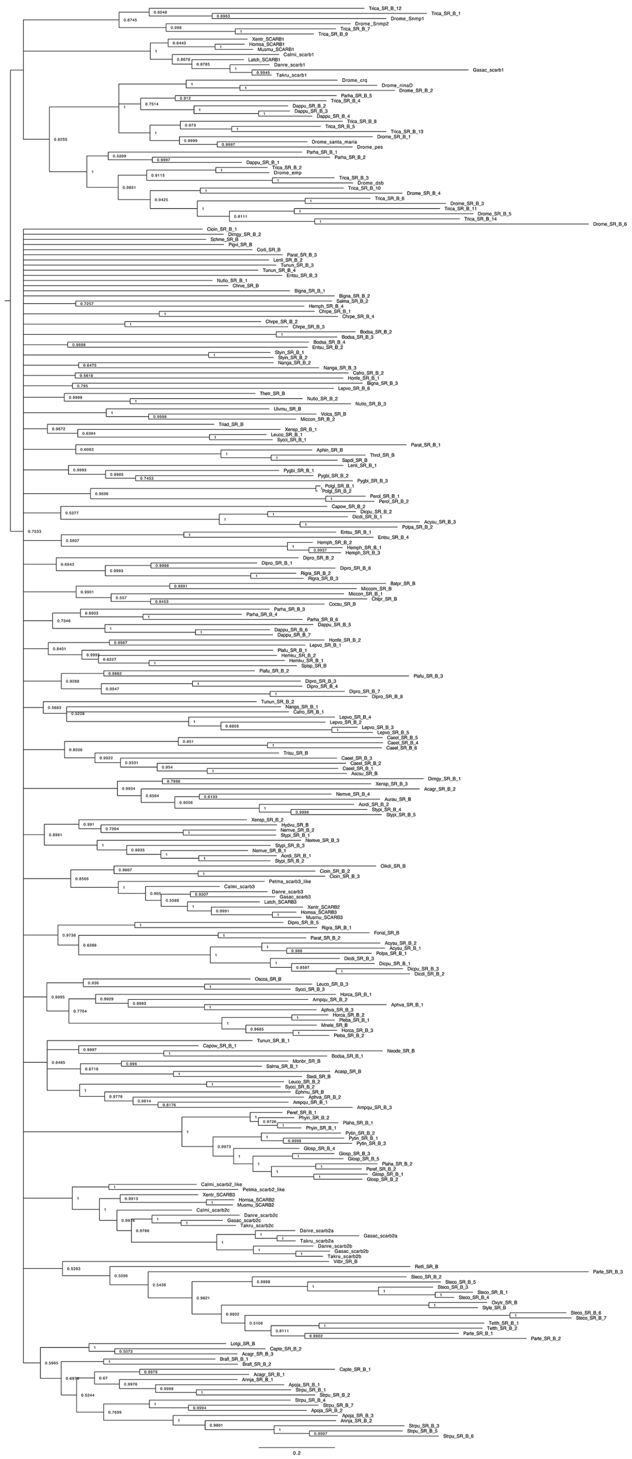

**Supplementary Figure 3: Bayesian phylogenetic analysis.** Best tree based on amino acid MAFFT alignment corresponding to 279 eukaryotic CD36 ectodomains. inferred via MrBayes v3.2.7 using the LG+G4 evolutionary model and two independent runs of 6 million generations. Labels indicate node bayesian posterior probability (BPP). Tip labels are protein names. Refer to Supplementary Table 2 for source genome or transcriptome, and sequence identifiers.

### Supplementary Figure 4

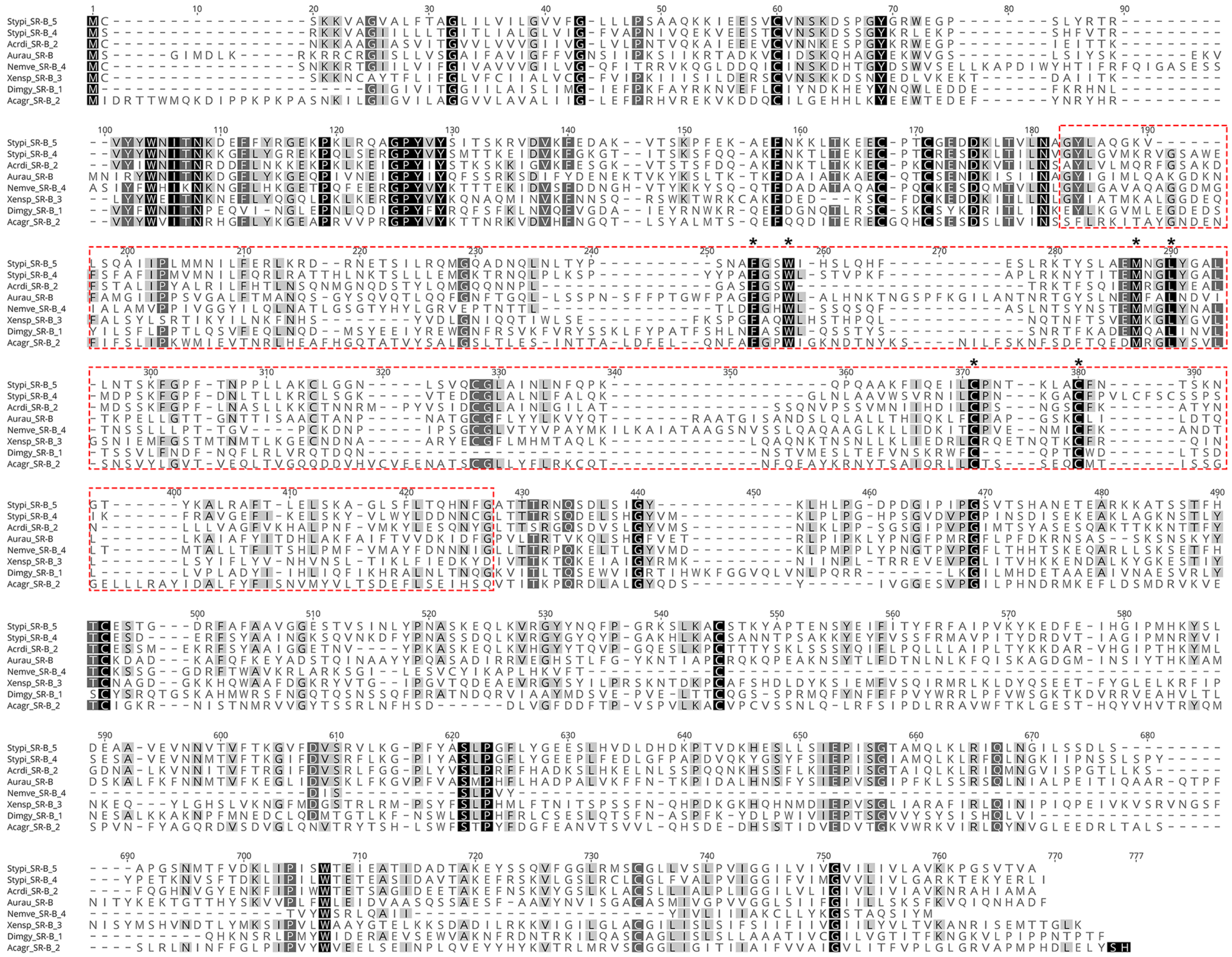


**Supplementary Figure 4. Alignment the eight cnidarian and lophotrochozoan CD36 ectodomains captured by RAxML as a bootstrap supported clade.** CD36 ectodomains from the eight cnidarian-lophotrochozoan sequences (Figure 2; Supplementary Figure 2) were aligned separately to investigate putative homology vs. the potential long-branch attraction. The red boxed region represents lineage-specific sequence expansions not present in any other metazoan SR-Bs (all other metazoan sequence lengths <500aa ). Aligned residues colored black 100% similarity, dark gray 80-99% similarity, light gray 60-79% similarity, and white <60% similarity. Pairwise identity across the highlighted sequence expansion region is 20.4%, with six invariant residues noted by asterisks (2.4% of sites).

**Supplementary Figure 5**

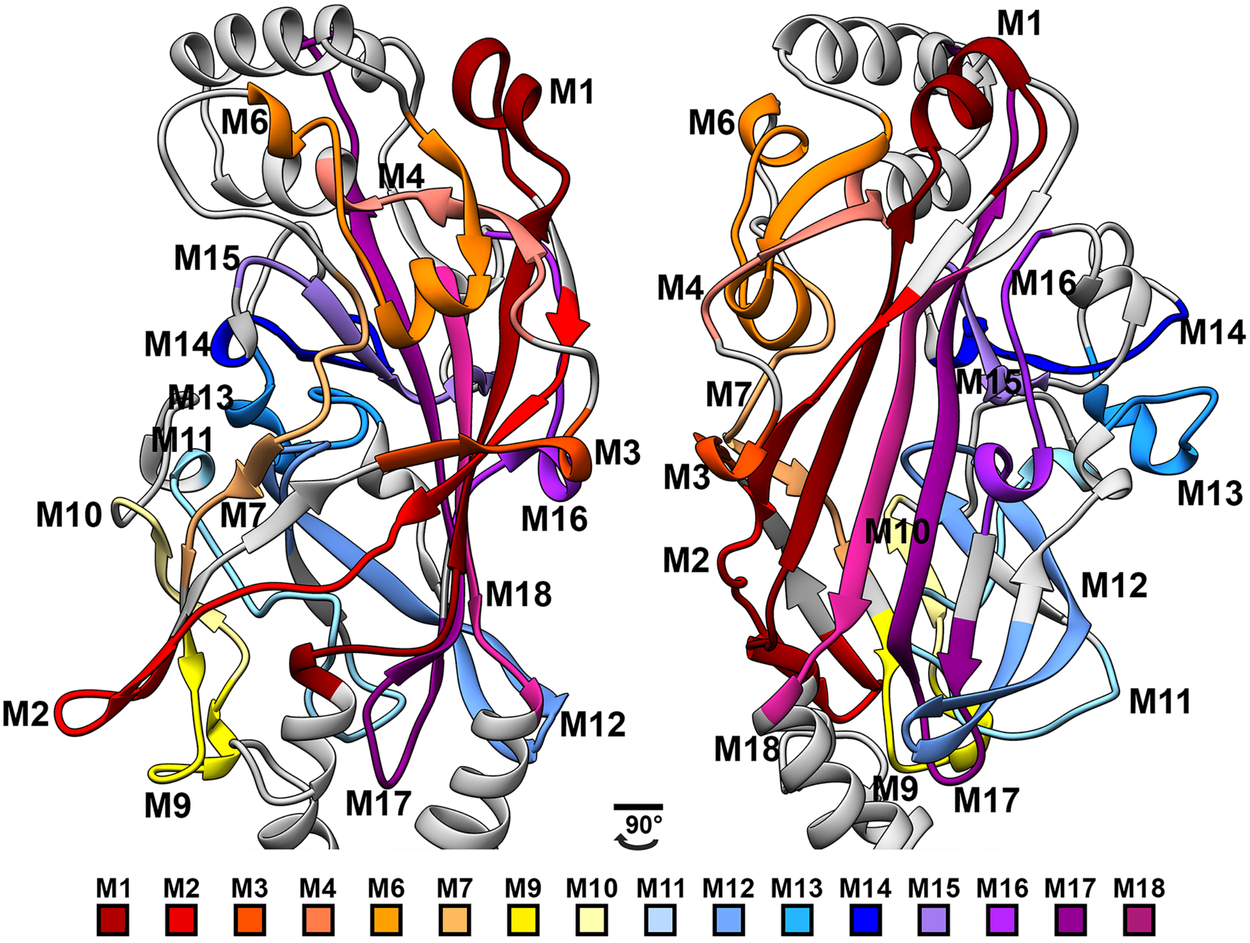


**Supplementary Figure 5: CD36 ectodomain amino acid motifs mapped to *H. sapiens* SCARB3 protein structure model.** STREME-identified CD36 ectodomain amino acid sequence motifs are color coded and mapped on corresponding residues of the human SCARB3 AlphaFold2 structure model. The model on the right is rotated 90°. Amino acid sequence motifs M5 and M8 were not detected in the human SCARB3 sequence.

### Supplementary Figure 6

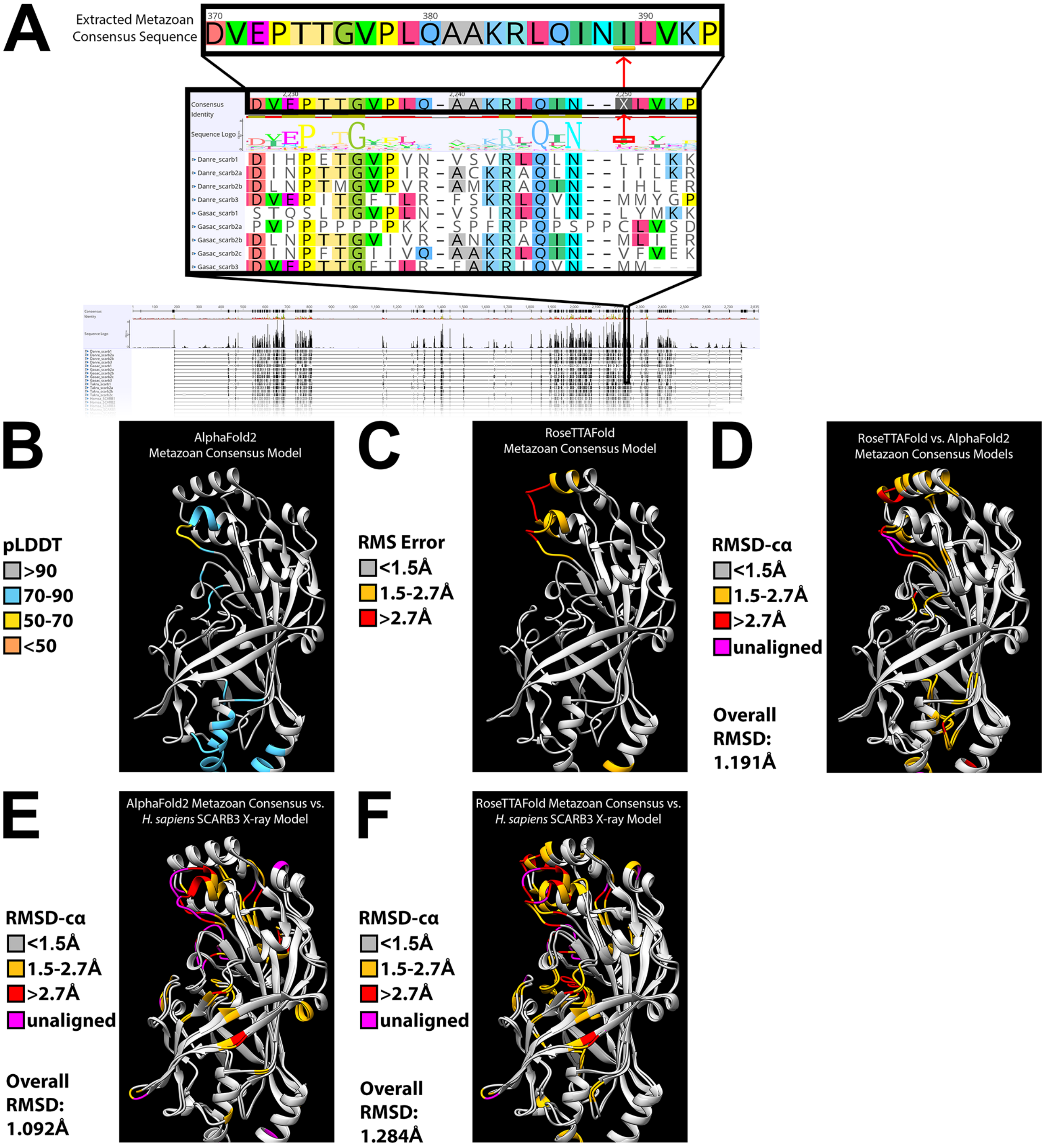

**Supplementary Figure 6: Metazoan consensus SR-B sequence and model generation.** We generated a generalized metazoan SR-B from a consensus sequence derived from the alignment of 143 metazoan sequences to illustrate the stereotypical metazoan CD36 ectodomain architecture, including the presence of a three alpha-helix bundle at the apex of the protein structure. **A)** Ambiguities in the consensus sequence were corrected by selecting the most common amino acid at that position, or by selecting the first amino acid given by the sequence logo. **B)** Residues in the AlphaFold2 model are colored by the predicted local distance difference test (pLDDT) confidence metric. Error ranges follow the "low" (<70), " medium" (70-90), and "high" (>90) pLDDT confidence ranges commonly applied to AlphaFold2 models (Jumper et al., 2021). **C)** Residues in the RoseTTAFold model are colored by the predicted root mean square (RMS) error, as calculated by RoseTTAFold in Ångstroms. Error ranges correspond to "high" (<1.5Å), "medium" (1.5-2.7Å), and "low" (>2.7Å) resolution commonly used to assess the quality of crystallographic protein models (Wlodawer et al., 2007). **D)** The metazoan consensus sequence protein predictions from AlphaFold2 and RoseTTAFold structurally aligned to each other with residues colored by RMSD-cα. **E)** The AlphaFold2 for the metazoan consensus sequence and *Homo sapiens* SCARB3 crystal structure structurally aligned to each other with residues colored by RMSD-cα. **F)** The RoseTTAFold for the metazoan consensus sequence and*Homo sapiens* SCARB3 crystal structure structurally aligned to each other with residues colored by RMSD-cα.

### Supplementary Figure 7

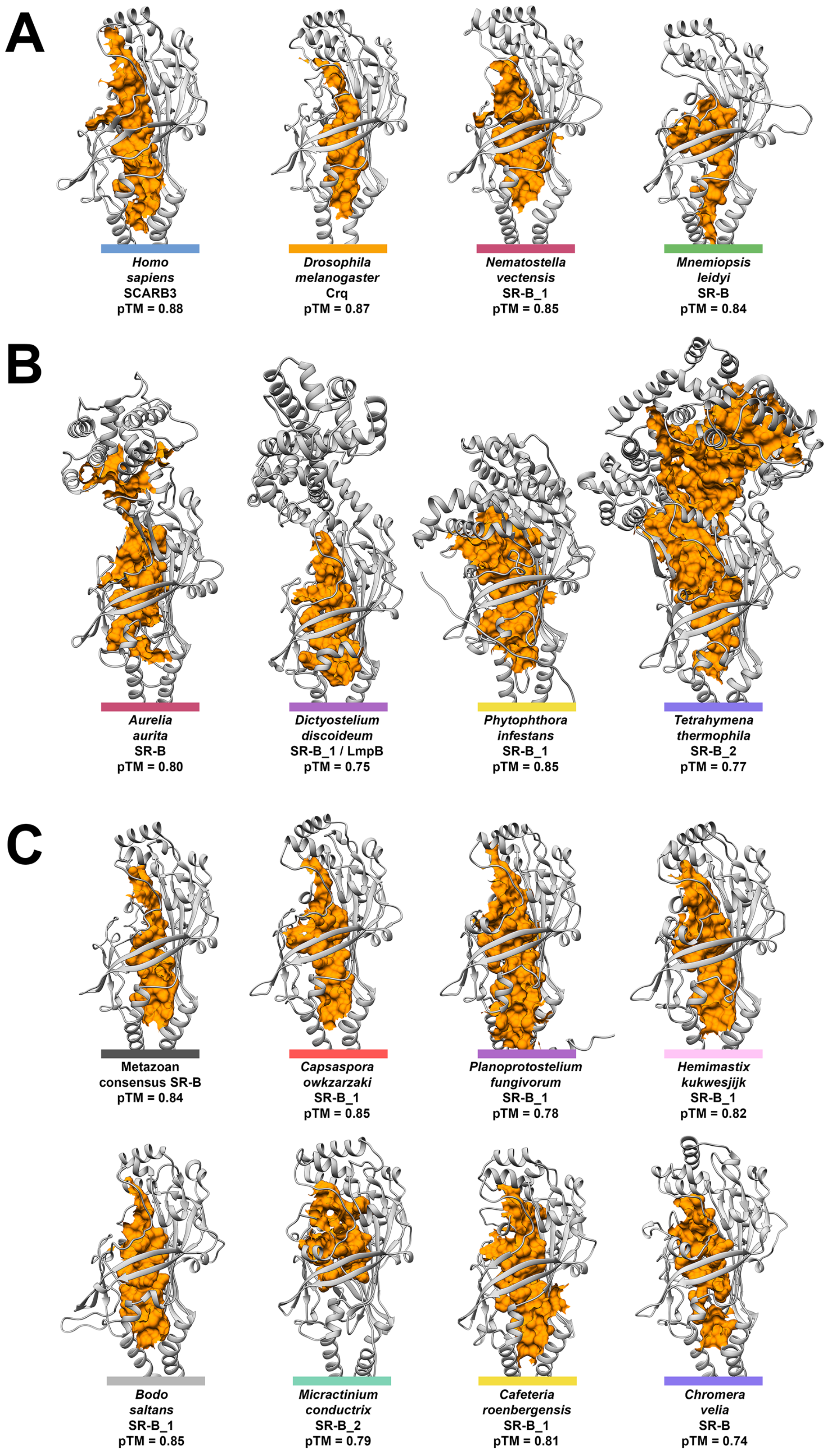

**Supplementary Figure 7: Intramolecular cavity and tunnel exemplars detected across eukaryotic SR-Bs.** The CASTp 3.0 web server (Tian et al., 2018) was used to predict the presence of intramolecular cavities in AlphaFold2-predicted SR-B protein structure models using a probe radius of 0.9Å (Chovancova et al., 2012). The surface of the largest cavity predicted by CASTp is indicated in orange with mouth atoms excluded. **A)** AlphaFold2 ribbon models from Figure 4 for metazoan SR-Bs with apex composed of a three-helix bundle. **B)** AlphaFold2 ribbon models from Figure 5 for eukaryotic SR-Bs with divergent apex helical bundle regions. **C)** AlphaFold2 ribbon models for non-metazoan SR-Bs with apex composed of a three-helix bundle.

### Supplementary Figure 8

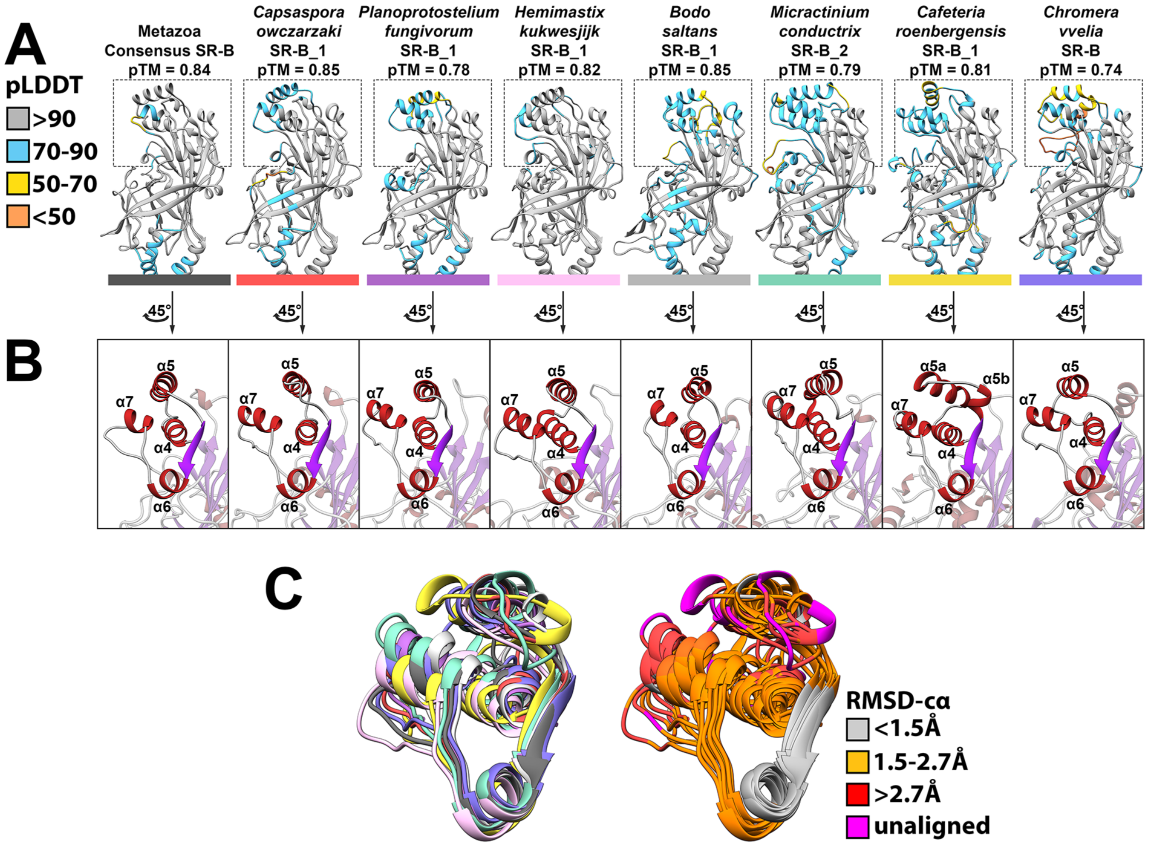


**Supplementary Figure 8: Structurally conserved three-helix bundle present across diverse eukaryotic groups.**

AlphaFold2 structure models for metazoan consensus SR-B and seven SR-Bs from diverse eukaryotes. **A)** Residues in each model are colored by the per-residue confidence metric, predicted local distance difference test (pLDDT). Error ranges follow the “low” (50-70), ”medium” (70-90), and “high” (>90) pLDDT confidence ranges commonly applied to AlphaFold2 models (Jumper et al., 2021). Residues <50 pLDDT not considered. The membrane-distal apex region is boxed. A predicted template-modeling score (pTM) is given for each model as a metric for global model confidence. Colored bars correspond to taxonomic groups established in Figure 1 (with the metazoan consensus SR-B being represented in black). **B)** α-helical regions of the CD36 ectodomain apex in each model highlighted and rotated 45° to depict structural similarity to the three-helix bundle present in most metazoan SR-Bs (represented here by the metazoan consensus SR-B). Predicted α-helices are colored red and beta strands are colored purple. α-helices in each model are labeled according to their spatial correspondence with mammalian SCARB3 α-helices. Helices α4, α5, and α7 comprise the ligand-interacting three-helix bundle of mammalian SR-Bs. The beta sheet and α6, that connect α5 to α7, correspond with Motif 6 (Figure 3). **C)** To the left, each model superimposed and colored by taxonomy (left). To the right, each model superimposed and colored by root mean square deviation per carbon atom (RMSD-cα). RMSD-cα values were binned as “high” (<1.5Å), “medium” (1.5-2.7Å), or “low” (>2.7Å) resolution (Wlodawer et al., 2007). Residues that fail to align are “unaligned” and colored magenta.

**Supplementary Table 1: Selected taxa, genomes and transcriptomes.** Reference information for each genome and transcriptome. Taxonomic affiliation, date of last access, download source, and associated citation where available. Colors correspond with the taxonomic scheme in Figure 1B. This table is related to Figure 1.

**Supplementary Table 2: CD36 search results.** Each CD36-containing protein in this study with sequence identifier/accession number, CD36 *hmmsearch* result e-value, full protein sequence length, and InterProScan protein architecture result. Only sequences containing a CD36 domain flanked by transmembrane domains were used for subsequent analyses. Colors correspond with the taxonomic scheme in Figure 1B. This table is related to Figure 1.

**Supplementary Table 3: Comparison between RAxML and Bayes CD36 gene trees.** Composition of SR-B clades united by nodes supported by BS (>80%) and BPP (>90%). This table is related to Figure 2.

**Supplementary Table 4: CD36 disulfide bridge conservation.** Six cysteine residues are associated with three disulfide bridges in mammalian SR-B homologs and referred to as disulfide bridge-1, disulfide bridge-2, and disulfide bridge-3. Homologous cysteine residue presence is indicated in green. Homologous cysteine residue absence is indicated in black. Each cysteine residue is labeled according to the corresponding position in the PFAM CD36 *hmm*model, human SCARB3, and the unfiltered MAFFT alignment of CD36 ectodomains (Supplementary File 3). Presence was reported if a cysteine residue in a sequence occupies the same position in the unfiltered MAFFT alignment of CD36 ectodomains as one of the mammalian bridge-forming cysteines. Cysteines from three saprolegnialean oomycete SR-Bs (Thrcl_SR-B, Aphin_SR-B, Sapdi_SR-B) were excluded as false positives, as the putative cysteine matches were found to belong to a region of lineage-specific sequence expansion upon visual inspection.

**Supplementary Table 5: STREME motifs.** The sequence match scores for each of the 18 amino acid motifs discovered by STREME are given for all 279 eukaryotic CD36 ectodomains used in this study. P-values and E-values are given for each motif, along with the motif's consensus sequence. Sequence scores have been transformed to a 0–100 scale by dividing each score by the maximum sequence score (75.54, representing the raw sequence score for M1 to Calmi_scarb2c). This table is related to Figure 3.

**Supplementary Table 6: pLDDT scores for AlphaFold2 models.** All 279 eukaryotic SR-Bs <1401 aa modeled by AlphaFold2, as well as a metazoan consensus SR-B occupy each column, with the pLDDT per-residue confidence metric generated by AlphaFold2 for each SR-B model occupying each row. The predicted template-modeling score (pTM) for each structure prediction is given as a metric of global accuracy. This table is related to Figures 4-6.

**Supplementary Table 7: Angstroms error estimates for RoseTTAFold models.** All 279 eukaryotic SR-Bs <1401 aa modeled by RoseTTAFold, as well as a metazoan consensus SR-B occupy each column, with the Ångstrom error estimate per amino acid position generated by RoseTTAFold for each SR-B model occupying each row. The estimated local distance difference test estimate (lDDT) for each structure is given as a metric of global accuracy.
